# High-resolution sweep metagenomics using fast probabilistic inference

**DOI:** 10.1101/332544

**Authors:** Tommi Mäklin, Teemu Kallonen, Sophia David, Christine J. Boinett, Ben Pascoe, Guillaume Méric, David M. Aanensen, Edward J. Feil, Stephen Baker, Julian Parkhill, Samuel K. Sheppard, Jukka Corander, Antti Honkela

**Affiliations:** Helsinki Institute for Information Technology HIIT, Department of Mathematics and Statistics, University of Helsinki, Finland; Department of Biostatistics, University of Oslo, Norway; Wellcome Sanger Institute, Hinxton, Cambridge, United Kingdom; Centre for Genomic Pathogen Surveillance, Wellcome Genome Campus, Hinxton, Cambridge, United Kingdom; Hospital for Tropical Diseases, Wellcome Trust Major Overseas Programme, Oxford University Clinical Research Unit, Ho Chi Minh City, Vietnam; Centre for Tropical Medicine and Global Health, Nuffield Department of Clinical Medicine, University of Oxford, Oxford, United Kingdom; The Milner Centre for Evolution, Department of Biology and Biochemistry, Bath University, Bath, United Kingdom; Department of Infectious Disease Epidemiology, Imperial College London, London, United Kingdom; Big Data Institute, Li Ka Shing Centre for Health Information and Discovery, Nuffield Department of Medicine, University of Oxford, Oxford, United Kingdom; Department of Public Health, University of Helsinki, Finland; Helsinki Institute for Information Technology HIIT, Department of Computer Science, University of Helsinki, Finland

## Abstract

Determining the composition of bacterial communities beyond the level of a genus or species is challenging because of the considerable overlap between genomes representing close relatives. Here, we present the mSWEEP method for identifying and estimating the relative abundances of bacterial lineages from plate sweeps of enrichment cultures. mSWEEP leverages biologically grouped sequence assembly databases, applying probabilistic modelling, and provides controls for false positive results. Using sequencing data from major pathogens, we demonstrate significant improvements in lineage quantification and detection accuracy. Our method facilitates investigating cultures comprising mixtures of bacteria, and opens up a new field of plate sweep metagenomics.

## Background

High-throughput sequencing technologies have enabled researchers to study bacterial populations in unprecedented detail using whole-genome sequencing (WGS) of pure individual bacterial colonies. Sequencing of individual isolates has provided insights into antimicrobial resistance and the complex ecology of the spread of antimicrobial resistant variants globally. The application of community profiling metagenomics, in which the 16S rRNA gene is sequenced from complex multi-species samples, can provide information about the composition and dynamics of highly diverse bacterial populations. However, the resolution of this approach is limited because assignment beyond the level of genus/species to individual variant is generally not possible due to insufficient nucleotide variation [1]. Whole-genome shotgun metagenomics delivers a much higher resolution than 16S rRNA sequencing [2] but widespread application is hindered by the cost associated with sequencing a sample to a sufficient depth to capture the diverse set of organisms that may be present in the sample [3].

Current methods for identifying bacteria from sequencing data usually focus on analysing predetermined single nucleotide variants (SNVs) and/or marker genes to capture the variation contained in a mixed colony [4–6], or on analysing isolated colonies. Many potential applications require an analysis of data from mixed colonies where methods developed for pure colonies are insufficient. Furthermore, whilst the SNV-based approach has been successful in studies of the history of the human population, focusing solely on SNVs inadequately captures the greater variability and different modalities of variation in bacterial genomes. Conversely, solely gene-based approaches can capture some of this while potentially losing finer detail. Therefore, we aimed to strike a balance between these two approaches by making use of a complete genome reference database.

Here, we have developed the mSWEEP method, which is designed to make efficient use of large collections of reference genomes that are available for numerous important human pathogens and other culturable bacterial species. mSWEEP combines clustering of the reference genomes into biologically relevant groups, fast pseudoalignment of reads to the references, fast and accurate probabilistic inference of the cluster abundances, and a method for controlling false positive detections.

Although applicable to any scenario where reference genomes for the sequenced bacteria are available, mSWEEP specifically enables a new kind of high-resolution analysis in plate sweep metagenomics, where a mixture of colonies is harvested from an enrichment culture by sweeping the whole plate in contrast to isolating a single colony. Plate sweep experiments fall between whole-genome sequencing of single colonies and culture-independent metagenomics by analysing the entire complexity of a community from a specific growth medium. As illustrated in our experiments, this setting is ideal for analysing samples representing populations of pathogenic bacteria, where the infecting species of primary interest have generally been previously encountered and sequenced frequently. By leveraging on existing high-resolution genomic pictures of pathogen populations, mSWEEP provides means to address a range of novel biological questions related to within-host variation, transmission, and the effect of ecological factors on the microbial diversity present in samples.

## Results

### Lineage identification

Abundance estimation with mSWEEP is performed in two phases: *reference preparation*, performed once for a given reference collection, and *analysis* of samples (Figure 1). Reference preparation consists of defining a reference sequence database and grouping the sequences according to biological criteria such as sequence types (ST), clonal complexes (CC), or by using a clustering algorithm for bacterial genomes. Grouping related reference sequences is essential in enabling identification of the taxonomic origin of each read [7] and enables abundance estimation when the sequencing reads originate from a sequence having no exact match in the reference database but which is represented by sequences from closely related organisms within the same group (typically bacterial lineage). Consequently, accuracy of the abundance estimates provided by mSWEEP is reliant on an extensive reference database and a biologically meaningful grouping.

**Figure 1.**
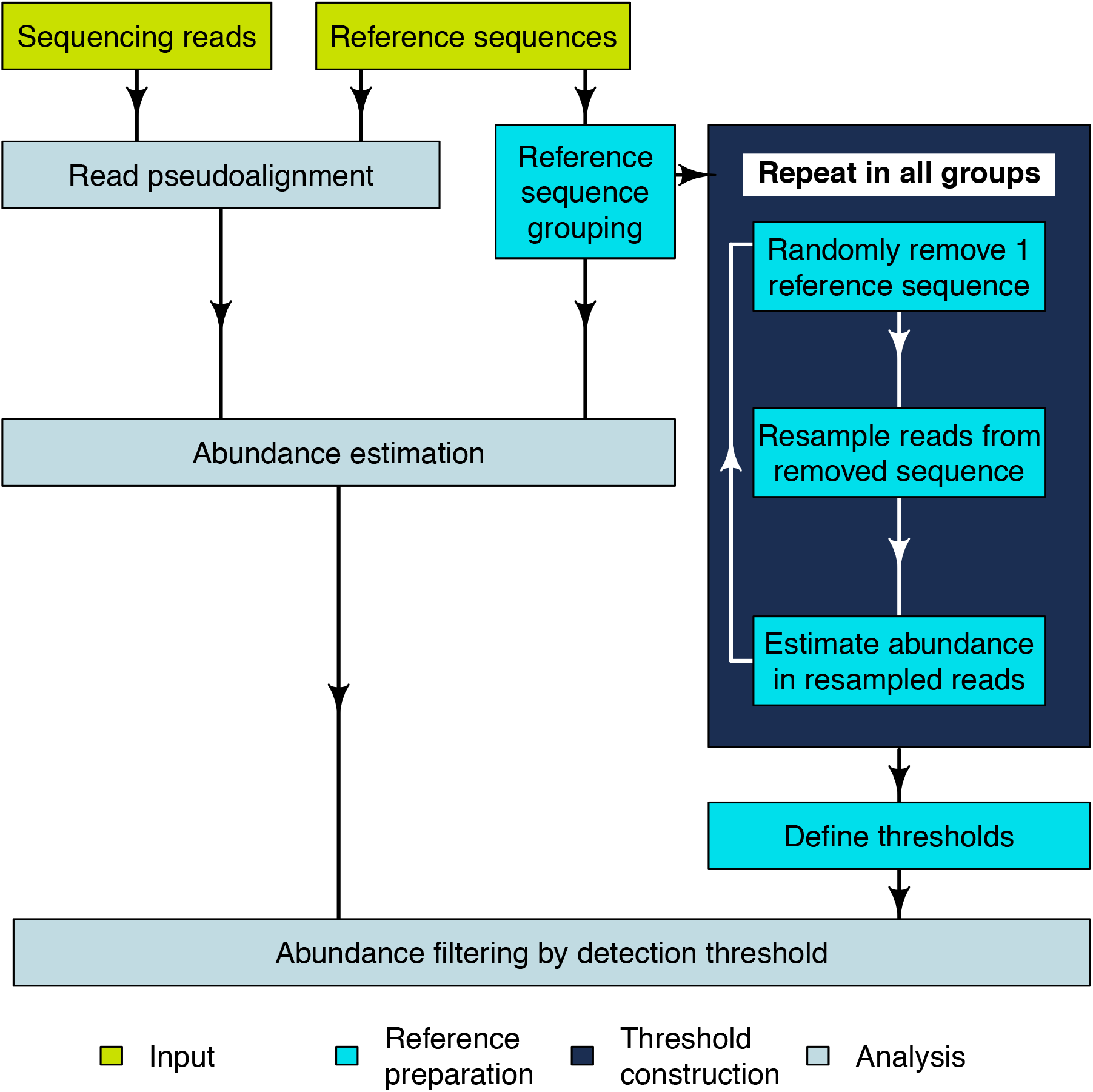
**Flowchart of the mSWEEP pipeline** describing a typical workflow for relative abundance estimation. The input part refers to the input data, reference preparation to the operations that need to be performed once per set of reference sequences, and analysis contains the steps run for every sample.

We constructed *detection thresholds* for the groups during the reference preparation from the reads used to assemble the reference sequences (Figure 1). We randomly chose one reference sequence at a time, removing it from the reference set, resampling the reads from the removed sequence, estimating abundances from the resampled reads, and repeating the process for all groups within the reference set. The detection threshold for a given group was determined by examining abundance estimates obtained when the group is not the true source and setting the threshold at the maximum false abundance estimate. This approach provides a statistical confidence score for estimates exceeding the detection thresholds, corresponding to the level of error deemed acceptable in abundance estimation from the resampled reads.

The first phase of analysis is pseudoaligning [8] sequencing reads to the reference sequences. Pseudoalignment produces binary *compatibility vectors* indicating which reference sequences a read pseudoaligns to. Based on the pseudoalignment count to each reference group, we defined the likelihood of a read originating from each of the groups. We assumed that 1) if multiple groups have the same total number of reference sequences, the group with a higher fraction of pseudoalignments is the more likely source for the read, and 2) the likelihood of the read to originate from a group is not dependant on the number of reference sequences in the group.

Basing the likelihood on the pseudoalignment compatibility vectors defines an extension of a probabilistic model that has previously been applied in RNA-sequencing [9, 10] and to bacterial data [7]. The extended model utilizes multiple reference sequences from each group as opposed to the previous attempts that rely on selecting a single, best-representative sequence from each of the groups [7]. Our model obtained the relative abundances of the reference groups by considering the generating process for a sample as a pooling of sequencing reads originating from the reference groups according to some unknown proportions, corresponding to a statistical mixture model. We fit the model and inferred the mixing proportions using variational inference [10].

### Assigning single-colony isolates to lineage

We compared the performance of mSWEEP against two existing methods capable of either strain or lineage identification: metakallisto [11] and BIB [7]. The main differences between the methods are that metakallisto attempts to identify individual strains based on all available sequences, BIB uses grouped reference sequences with a single representative sequence from each group to assign abundances to the groups, and mSWEEP identifies the presence of groups by using grouped reference sequences with all the available sequence as representatives.

As the reference data, we used bacterial sequence assemblies from four studies [12–15] augmented by single representative sequences from 27 species; a total of 3815 reference sequences. We grouped the sequences in either clonal complexes, lineages identified with the BAPS clustering algorithm [16], or on the species-level. We removed 504 sequences from all groups represented by more than one sequence to create a dataset where the true group is known but the true sequence is not available to the method pipeline (Table 1). In addition to the test data described in Table 1, we referred to a study sequencing 61 *K. pneumoniae* isolates from Thailand [17] to assess the accuracy of all methods when the reference sequences and the test samples were not obtained from the same source.

**Table 1.**
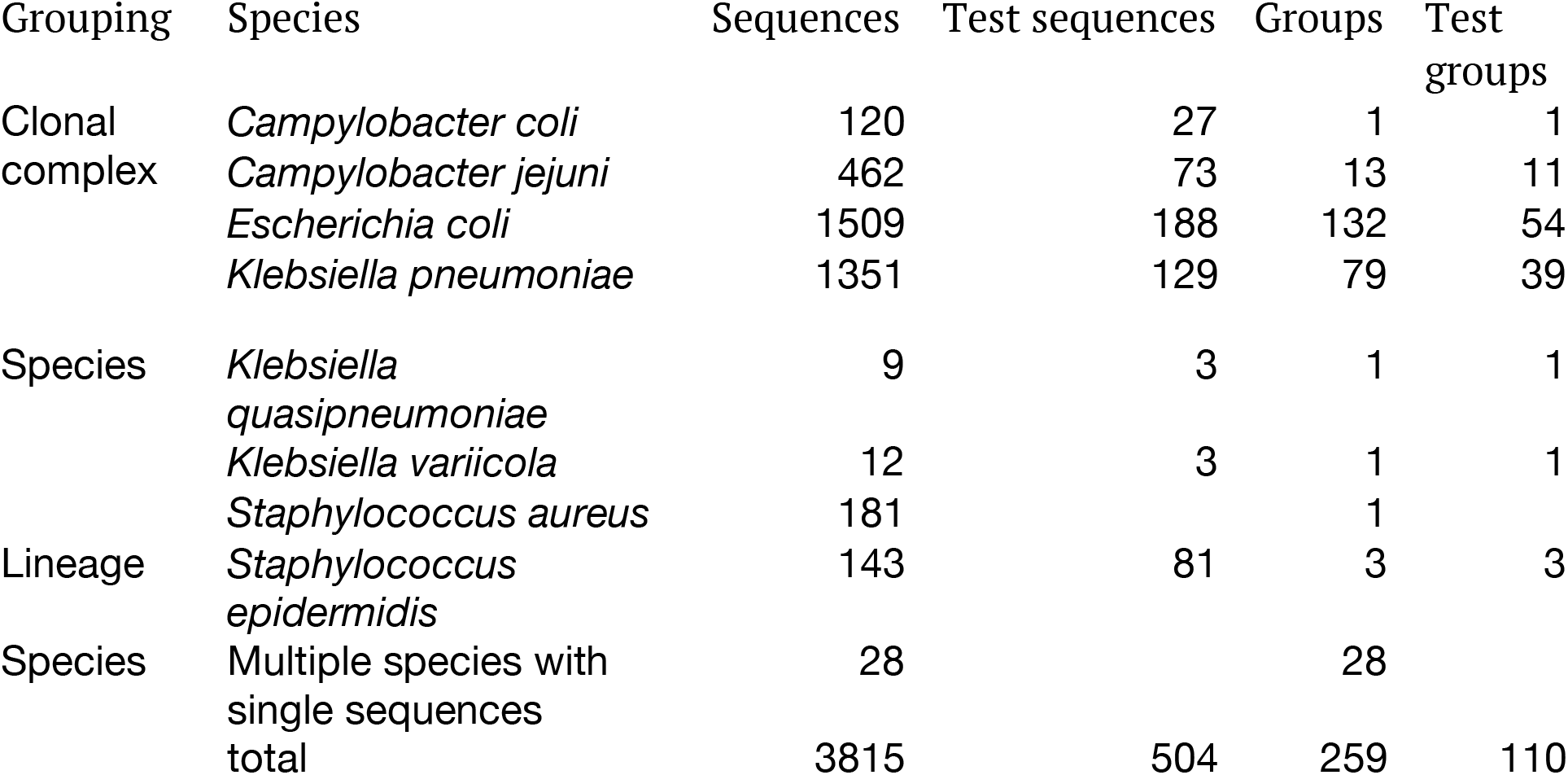
**Reference data** used to perform the analyses and to evaluate the performance of mSWEEP, metakallisto, and BIB. Clonal complexes are defined as either single-locus variants from the central sequence type (*Campylobacter jejuni, Campylobacter coli)* or double-locus variants (*Klebsiella pneumoniae* and *Escherichia coli*). The *Staphylococcus epidermidis* lineages were identified in the original study with the BAPS clustering algorithm.

mSWEEP significantly outperformed BIB and metakallisto in cases measuring accuracy of abundance estimates in the true group (Figure 2; p < 10^−9^, in all comparisons, Wilcoxon signed-rank test; median error in all estimates for mSWEEP was 0.00003, for BIB 0.23, and for metakallisto 0.54). When measured by highest estimates in the incorrect groups, mSWEEP outperformed the other two methods in all cases except the *S. epidermidis* 11-group clustering and the *K. pneumoniae* out-of-reference samples (Figure 2; p < 0.0012, Wilcoxon signed-rank test; median error in all estimates for mSWEEP was 0.000002, for BIB 0.05, and for metakallisto 0.01). In these two latter cases, mSWEEP and metakallisto performed similarly (Figure 2; p > 0.10 when testing for the difference in accuracy in either direction, Wilcoxon signed-rank test). Since metakallisto attempts to identify strains rather than lineages, the observed behaviour is likely a result of the majority of the abundance estimates being spread across strains belonging to the true lineage.

**Figure 2.**
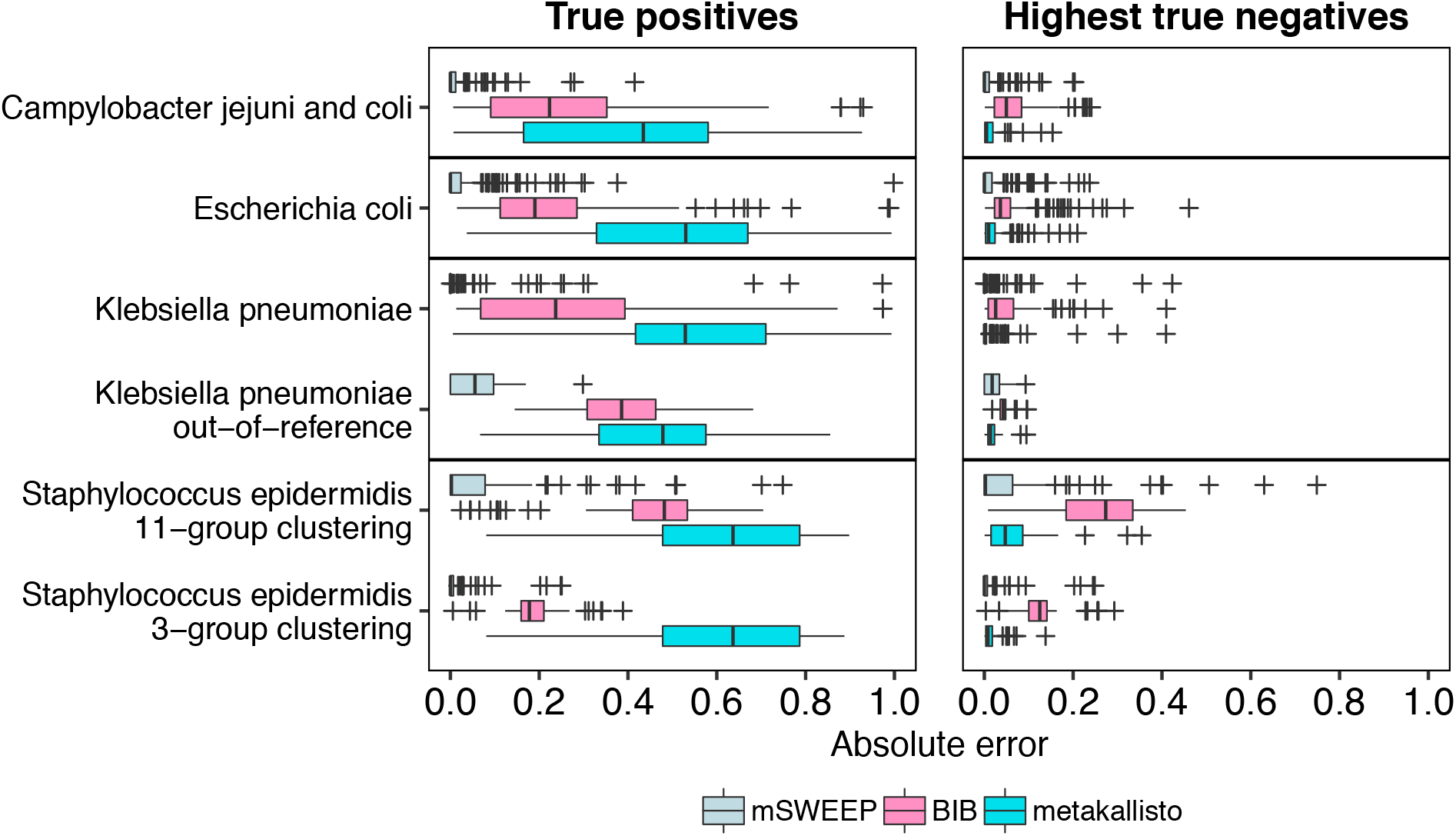
**Error of abundance estimates in single-colony isolates** (lower is better). True positives represent the relative abundance estimates in the true lineage (mSWEEP and BIB) or the highest estimate for strains of the lineage (metakallisto). Highest true negatives contain the highest estimate in an incorrect lineage (mSWEEP and BIB) or the highest estimate for a strain from an incorrect lineage (metakallisto). The absolute error is deviation from an abundance of one (True positives) or zero (Highest true negatives).

We additionally compared mSWEEP and BIB by measuring accuracy in classification based on assigning the samples to the lineage with the highest abundance estimate. With this criterion, both methods correctly identified the true clonal complex in all 100 *C. jejuni* and *C. coli* isolates, and in all 81 *S. epidermidis* isolates when using the 3-cluster grouping. In the 11-cluster *S. epidermidis* grouping, mSWEEP correctly identified the true lineage in 78 and BIB in 80 samples. In the 188 *E. coli* and 129 *K. pneumoniae* isolates, mSWEEP identified the lineage correctly in 187 and 126 samples, while BIB correctly identified 184 and 117. The *K. pneumoniae* and *E. coli* isolates that were misidentified by mSWEEP likely contain a sequence type that is missing from the reference, or are mixtures of *K. pneumoniae* and *E. coli* lineages (Supplementary Figures 1a and 1b). Out of the last 61 out-of-reference *K. pneumoniae* samples, mSWEEP identified the true origin in all 61 isolates and BIB in 53.

The least accurate estimates for all methods (measured by the true positives and highest true negatives) were obtained for the 81 *S. epidermidis* isolates when using the second level of the hierarchical BAPS clustering with 11 groups (Figure 2), where none of the three methods reached the level of accuracy observed in the other cases. These inaccuracies are explained by the comparably small reference for the *S. epidermidis* population (Table 1), which does not exhibit a clear cluster structure (Supplementary Figure 2a) beyond the coarsest BAPS clustering into three groups. The lack of structure causes the abundance estimates to spread across the new groups defined within each of the three top-level clusters (Supplementary Figure 1c).

We further examined the grouping of the reference sequences by producing t-SNE plots of *31*-mer distances between the reference sequences including the test isolates (Figure 3, Supplementary Figures 2a-c). The *C. jejuni* and *C. coli*, *E. coli*, and *K. pneumoniae* references conform to the clonal complex grouping while the *S. epidermidis* population only conforms to the coarsest 3-group BAPS clustering. The t-SNE plots correctly place the assemblies into the true groups but the method does not preserve the distances between the points or the clusters [18] and is unsuited to analysing mixed isolate data.

**Figure 3.**
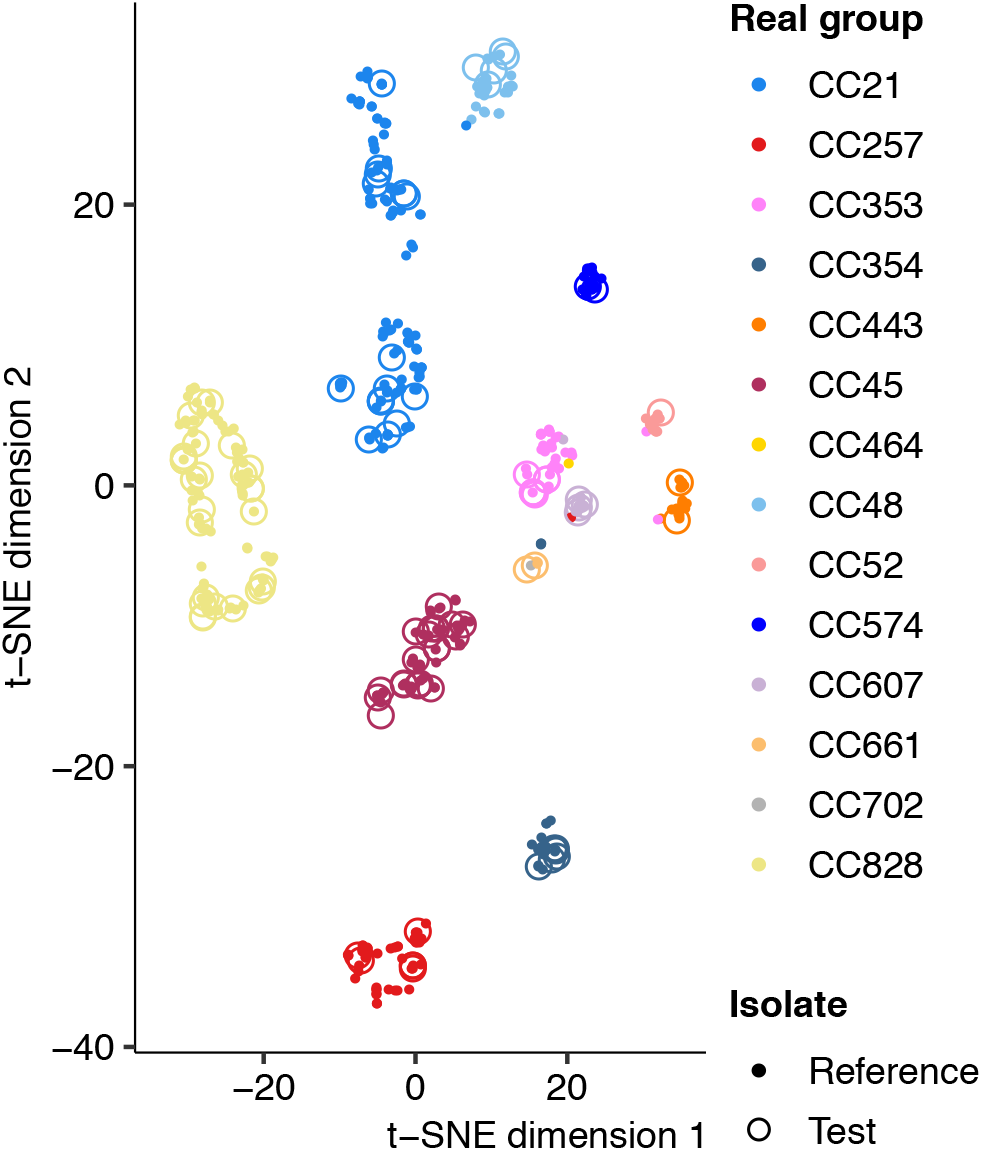
***C. jejuni* and *C. coli* reference 31-mer embedding.** t-SNE embedding of the 31-mer distances between the reference isolates shows that the reference population conforms relatively well to the clonal complex grouping. The test cases, indicated by circles, are all correctly identified by mSWEEP and t-SNE also places them within or near the true source group.

Processing the 504 single-colony isolates with mSWEEP took an average of 23 minutes and 50 seconds per sample, metakallisto an average of 24 minutes and 42 seconds, and BIB an average of 143 minutes and 46 seconds per sample using the same reference data. mSWEEP used a maximum of 79.5Gb RAM (maximum of 24.6Gb counting only the abundance estimation step), metakallisto 108.1Gb, and BIB 31.5Gb. Resource usage was obtained by running each sample separately with eight processor cores against the full test reference of 3311 sequences in 259 groups (mSWEEP and metakallisto) or a representative sequence from each group (BIB). Both mSWEEP and metakallisto could be sped up by running the pseudoalignment in kallisto’s batch mode (processing multiple samples simultaneously), reducing the need to write and read from the hard disk at the cost of increased memory usage.

### Quantifying synthetic mixtures of single-colony reads

We investigated the performance of mSWEEP in quantifying samples containing multiple lineages of bacteria from the same species by synthetically mixing reads from the single-colony samples. Each mixture sample was set to contain a total of one million reads from three single-colony samples from three lineages, with randomly assigned proportions from the set (0.20, 0.30, 0.50). We used a balanced incomplete block design to ensure that all lineages appear in at least 13 mixture samples, and each single-colony isolate appears at least once, producing 161 *C. jejuni* and *C. coli*, 477 *E. coli*, 584 *K. pneumoniae*, and 100 *S. epidermidis* synthetic mixture samples in total.

Compared to abundance estimates from the single-colony samples, estimates obtained from the synthetic mixture samples show that the presence of sequencing reads from multiple lineages in a synthetic mixture results in an error distribution resembling the one observed in the single-colony samples (Figure 4, Supplementary Figure 3). Estimates from the synthetic *S. epidermidis* mixture samples using the 11-group split produce an error distribution that differs from the single-colony error distribution more than that observed with the other groupings.

Comparing the empirical distributions of the errors from the synthetic mixtures and the single-colony isolates (Supplementary Figures 3 and 4) shows that for estimates exceeding a threshold of 0.016, the accuracy of estimates from the mixture samples stochastically dominates the accuracy observed in the single-colony samples, except in the *S. epidermidis* 11-cluster case where stochastic dominance is observed only above a threshold of 0.17. Stochastic dominance establishes a partial ordering between two random variables and, in this case, implies that estimates from the mixture samples are more accurate (in a probabilistic sense) than estimates from the single-colony samples when the estimates are large enough. In the *S. epidermidis* 11-cluster case we do not establish the mixture estimates as more accurate since the distribution (Supplementary Figure 3) and the observed threshold differ considerably from the other cases.

**Figure 4.**
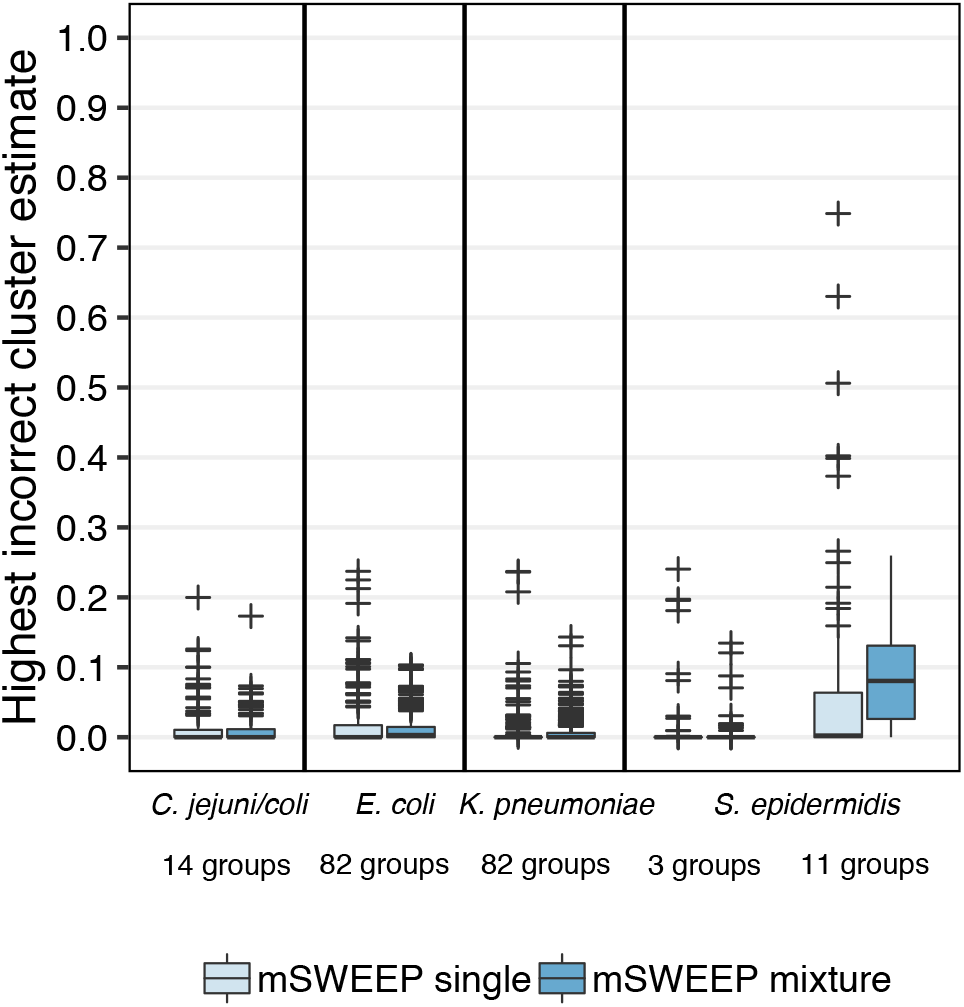
**False positives in single-colony samples versus synthetic mixtures.** Abundance estimates from synthetic mixtures of three lineages do not result in higher number of false positive estimates when compared to estimates from the single-colony samples, as measured by the largest estimate for a lineage that does not contribute any sequencing reads. The only exception is the *S. epidermidis* 11-cluster case which is not accurately identified in neither the synthetic mixtures nor the single-colony samples.

The results indicate that above this relatively low background noise level of 0.016, quantifying mixture samples is not expected to produce more false positive results than would be obtained from single-colony samples. This justifies simplifying the problem of determining the detection thresholds accompanying mSWEEP, which provide a threshold for reliable detection of the reference groups in mixture samples, to determining the thresholds based on the single-colony isolates. Due to the requirement that the abundance estimates must be large enough for this assumption to hold, we incorporate the threshold observed in comparing the estimates into the detection thresholds by using it as the minimum threshold regardless of the results from the resampling procedure.

### Results from plate sweeps

We applied the mSWEEP pipeline to three datasets containing multiple lineages of the same species: 116 samples from *C. coli* and *C. jejuni*, 96 paired samples from *E. coli*, and 179 samples from *K. pneumoniae*. The *E. coli* samples were obtained from MacConkey plate sweeps from a cohort study of 48 Vietnamese children during a diarrhoeal episode (48 samples), and when healthy (48 samples), purposefully expecting multiple lineages in each sweep. Conversely, the *C. coli*/*C. jejuni* and *K. pneumoniae* datasets were presumed pure cultures but flagged during downstream analysis as mixed. In all three experiments, we applied the detection threshold procedure (described in more detail in the Methods section) to filter the resulting abundance estimates. We used two thresholds, corresponding to confidence scores of 0.99 and 0.90, from now on referred to as filtering by 0.99 or 0.90 confidence thresholds.

### Population structure of commensal *Escherichia coli* from Vietnamese children

The most abundant sequence type complex identified in over half the samples (diarrhoeal and control samples) was CC10 (Figure 5, Supplementary Table 1). Notably, 95% of the samples (46/48 Diarrheal and 45/48 Healthy) harboured multiple antimicrobial resistance genes (identified using the ARIBA software [19]) that belonged to three or more classes of drugs (Supplementary Figure 5), which we defined as multi-drug resistance [20] (MDR. One sample was found to contain the plasmid associated resistance gene MCR-1, which confers resistance to colistin, a last line antimicrobial drug [21]. We found no significant difference in the antimicrobial resistance gene profile between the healthy and diarrhoeal samples (Two tailed, Fisher’s exact test p=0.5). Supplementary Table 2 details how many samples harboured antimicrobial resistance genes in each antimicrobial drug class; the full antimicrobial resistance gene data can be found in Supplementary Table 3.

**Figure 5.**
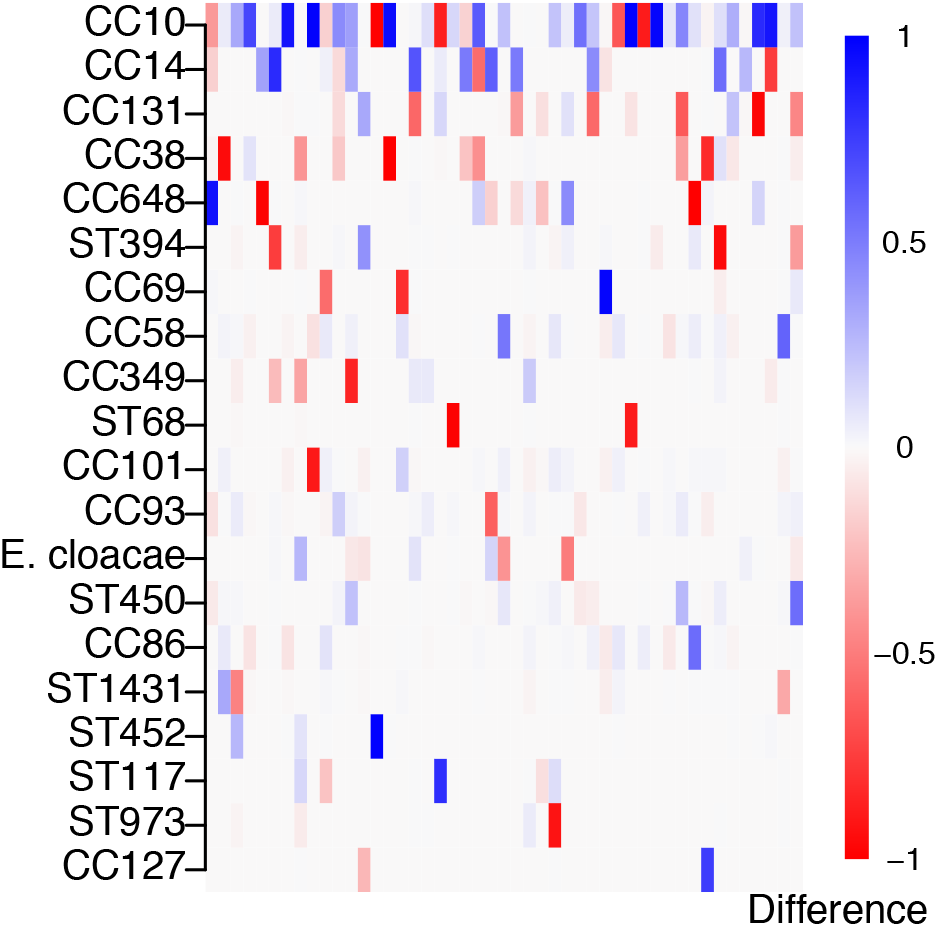
**Difference in *Escherichia coli* clonal complex (CC) and sequence type (ST) lineage abundances during and after diarrhoea.** The plot displays the differences in unfiltered relative abundance estimates before and after treatment in 20 most common (defined by the sum of relative abundances; blue denotes increase, red decrease) *E. coli* reference lineages or other species across all 47 paired samples represented in the columns.

We additionally examined differences between the community composition in the healthy and diarrhoeal samples based on both the distribution of the relative abundance estimates (alpha diversity), and changes in the identified strains (beta diversity). The alpha diversity, measured by Shannon entropy (Supplementary Figure 6), showed no significant differences between the two paired samples (p > 0.90, Wilcoxon signed-rank test; median Shannon entropy in diarrhoeal samples was 0.60, and in healthy samples 0.59). However, we found significant shifts in lineage composition (see Figure 5) when comparing the beta diversity, measured by Bray-Curtis dissimilarity, between the two samples (p < 0.005, multivariate-ANOVA). Tests were performed on relative abundance estimates filtered by both 0.99 and 0.90 confidence thresholds.

### Co-occurrence patterns in *Campylobacter* lineages

The network diagram (Figure 6a) shows ST-clonal complex (CC) (nodes) of the isolate genomes with the thickness of edges representing the number of times that isolates from these CCs are found together in a single plate sweep sample, and the size of the node the total number of observations. The overall amount of co-occurrence between CCs (Figure 6b) provides basic information about the frequency that CCs are found together in natural populations. *C. jejuni* CCs 45, 661, 607, 353, 48, and *C. coli* CC828 are all found in samples with 4 or more other CCs and there is evidence that isolates from some CC’s cohabit with other species including *Campylobacter lari* and *Bacillus subtilis*. While the sample set in this study was deliberately selected to include mixed isolate samples, quantifying the co-occurrence of species and lineages can provide information about different ecologies or lineage interactions, particularly when CCs are known to have varied sources, such as different hosts.

**Figure 6.**
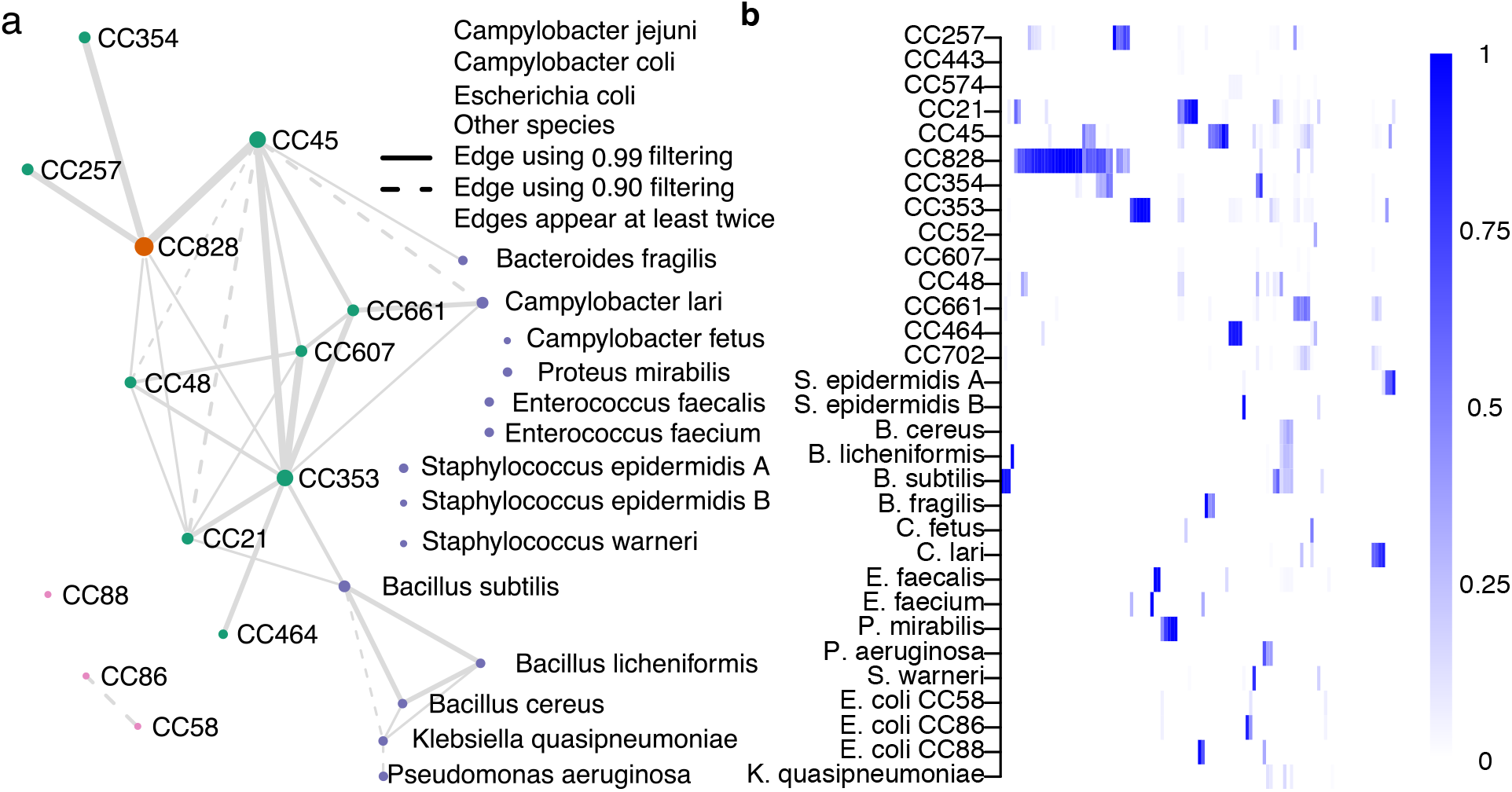
***C. jejuni* and *C. coli* clonal complex (CC) coexistence in 116 samples.** The coexistence network in panel a was constructed from relative abundance estimates filtered by detection thresholds constructed using a confidence score of either 0.90 (dashed edges) or 0.99 (solid edges). An edge between two groups represents coexistence in at least two samples with the chosen threshold. Edge size is proportional to the number of times the joined nodes were observed together, and node size to the total times the group was detected. Panel b visualizes the unfiltered relative abundance estimates in the same reference groups (rows) as in panel a, across 116 samples (columns).

There is some preliminary evidence that common clinical lineages CC45 and CC21 are rarely found together in a single sample (plate sweep) while other lineages, such as the chicken associated CC353, are frequently isolated from samples containing multiple strains. From an evolutionary perspective, it is unlikely that closely related strains can stably occupy identical niches because competition would be expected to lead one to prevail. The results demonstrate co-occurrence of strains within individual host animals and multi-strain infections in humans and provide information about the complex ecology of co-occurring interacting species that leads to the observed community structure in a given sample.

### Multi-drug resistant *Klebsiella pneumoniae* coexist with other lineages

The coexistence network (Figure 7) and the sample-lineage heatmap (Supplementary Figure 7) for the 179 human clinical samples of *K. pneumoniae* demonstrates common co-occurrence of *K. pneumoniae* with a wide variety of *E. coli* lineages, as well as occasional co-occurrence with *Acinetobacter baumanii* and other species. Both *E. coli* and *A. baumanii* grow on the media used for culture of *K. pneumoniae*. Clonal complexes of *K. pneumoniae* centred on sequence types associated with high levels of multi-drug resistance (e.g. ST258, ST147 and ST101) were frequently observed co-existing with a variety of other *K. pneumoniae* lineages as well as with each other, and with other important Gram-negative pathogens. Developing a deeper understanding of these community structures and interactions will be critical for monitoring horizontal transfer of antimicrobial resistance genes between taxa.

**Figure 7.**
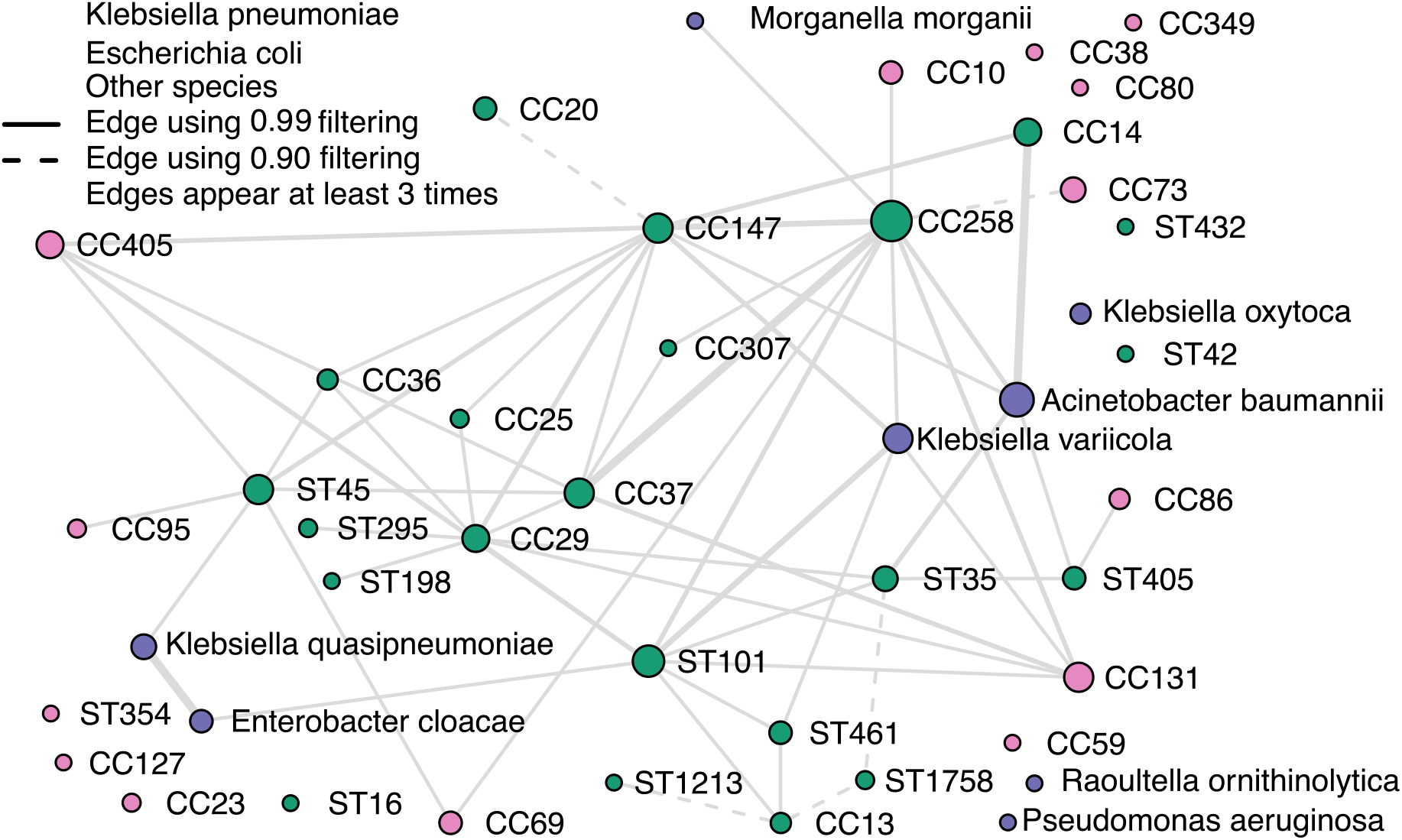
***K. pneumoniae* lineage coexistence in 179 mixture samples.** The coexistence network was constructed from relative abundance estimates remaining after filtering by the detection thresholds. Visible edges denote coexistence in at least three samples. Dashed edges represent coexistence when using detection thresholds corresponding to the 0.90 confidence score, and solid edges using the 0.99 detection threshold. Node sizes are proportional to the number of times the lineage was observed; edge sizes to the number of times coexistence was established.

## Discussion

Metagenomics using high-throughput sequencing has become a common approach when investigating the bacterial composition of different environments or changes introduced by intervention, such as in the human gut microbiome. In most epidemiological applications, the relevant target organisms are culturable using established media which offers a clear advantage to obtaining high sequencing depths in a cost-effective manner. We developed the mSWEEP pipeline to enable high-resolution inference of the lineages present in plate sweeps of enrichment cultures. mSWEEP can be used to infer the detailed population structure of a single species, or the diverse populations of bacteria typically encountered in clinical and public health settings where standard culturing media is routinely used to isolate epidemiologically relevant organisms. This pipeline also estimates the relative abundance of lineages and reliability cut-offs. mSWEEP was designed to have a minimal execution time using the latest advances in RNA-seq analysis and its maximum memory footprint is determined by the pseudoalignment algorithm. We demonstrated significant improvements in accuracy over the previous state-of-the-art method in our experiments.

mSWEEP provides considerable power for improving our understanding of infection by recovering a true representation of bacteria in a complex sample. For example, genotyping studies have shown that *C. jejuni* and *C. coli* colonizing the primary host (birds and mammals) form clusters of related isolates that are host-associated [22], which can be used to identify the reservoir for human infection [23]. However, multiple organisms can be isolated from the same sources [24, 25]. The co-occurrence of different organisms could be a snapshot in time of a wider process of lineage succession [26] in which the resident microbiota might resist new colonizations or be displaced by recently acquired bacteria [27, 28]. Further, we suggest we may be able to infer complex interactions between organisms that occupy different microniches [29] and are not in direct competition [30, 31] by analysing their co-occurrence. Therefore, this approach provides a means to investigate the nature of polymicrobial infections to improve our understanding of the spread of a specific organism between hosts and transmission to humans in addition to enabling characterization of physical and temporal variation in the distribution of lineages among multi-strain samples.

Because of limitations in the initial culture and DNA isolation processes, we can only infer relative (not absolute) abundances. However, this is not a significant limitation as the absolute abundances of target organisms are also subject to large biological and technical variation. Memory requirements for large reference collections or simultaneous analysis of multiple samples necessitate a dedicated computer cluster to run the analysis pipeline, but even for very large reference collections the resource usage is still at the level available at most bioinformatics centres. Alternatively, the reference sequences can be modified to include only the directly relevant species, which makes the method widely applicable to biologists. As with any method intended to identify sequence variation, the target species need to be relatively well known to allow building of sufficiently informative reference databases. Similarly, to allow for sensible and easily interpretable inferences, the biological clustering of the reference database should be based on well-established biological entities, such as multi-locus sequence types (STs) or clonal complexes (CCs) which are frequently employed as labels of lineages. As a by-product, mSWEEP can also be used to estimate the quality of the reference collection and to discard contaminated or mis-identified genomes.

Strain identification from metagenomic data has been recently suggested by the StrainPhlAn method [32]. mSWEEP, and similar methods, are complementary to StrainPhlAn as these methods analyse similar data but from different directions. mSWEEP assigns strains present in the sample to biologically established genetically separated lineages and estimates the relative abundance of these, whereas StrainPhlAn infers SNPs and phylogenetic relations within the whole sample. Given the flexibility and generality of the mSWEEP approach, we anticipate this method will pave the way for numerous novel applications of plate sweep metagenomics in many fields of microbiology.

## Conclusions

mSWEEP represents a novel means to quantify the composition of bacterial communities beyond the resolution offered by bacterial identification methods based on 16S ribosomal RNA gene sequencing or whole-genome shotgun metagenomics. We have demonstrated significant improvements in accuracy over similar methods, and novel co-existence analyses using plate sweeps of enrichment cultures of the human pathogens *Campylobacter jejuni*, *Campylobacter coli*, *Escherichia coli, Klebsiella pneumoniae* and *Staphylococcus epidermidis*. We expect that mSWEEP will find use in similar studies of bacterial pathogens, where high-resolution inference is required, ample reference collections for the species of interest are available, and the plate sweep metagenomic approach can be applied in-depth at a fraction of the current cost of single-colony sequencing.

## Methods

### Reference construction

The reference sequences (Table 1, Supplementary Table 4) are the genomic assemblies of a number of strains or species that represent the organisms of interest in a sample. We used a collection of assemblies from four studies [12–15] augmented with the genomes of a representative strain from 27 species that were identified in the real mixture data by MetaPhlAn [33].

### Grouping the reference sequences

We used the multilocus sequence types of the *C. coli, C. jejuni, E. coli* and *K. pneumoniae* reference sequences to group them into clonal complexes defined by the allelic profile of a central sequence type, and all other sequence types that vary in at most a single MLST locus (*C. coli* and *C. jejuni*) or in at most two loci (*E. coli* and *K. pneumoniae*). The *K. pneumoniae* reference contained sequences belonging to *K. variicola*, *K. quasipneumoniae*, and *K. quasivariicola* which we assigned to three groups defined by the three species. We similarly treated the 181 *S. aureus* contained in the *S. epidermidis* study as a single group, and split the 143 *S. epidermidis* sequences using the first and second levels of the hierarchical clustering produced by the hierBAPS [16] software (version 6.0). The complete grouping is provided in Supplementary Table 4.

### Pseudoalignment

We used kallisto [8] (version 0.45) with default settings to perform pseudoalignment. Pseudoalignment produces binary compatibility vectors which indicate whether the read pseudoaligns to a reference sequence or not. In our model, we sum the pseudoalignment counts within each reference group and thus consider the observations of the reads *r*_*n*_ = (*r*_*n, 1*_,…,**r**_*n, K*_), *n* = *1,…,N* as containing only the information about the number of pseudoalignments *r*_*n,k*_ within each of the *K* groups.

### Abundance estimation model

We assume that the reads *r*_*n*_ are conditionally independent given the mixing proportions of the groups *θ* = (*θ*_1_, …, *θ*_*K*_), and augment the model with the latent indicator variables *I* = *I*_1_, …, *I*_*N*_ which denote the true source group of each read. The joint distribution of the collection of reads *R* = *r*_1_, …, *r*_*N*_, the indicator variables *I* = *I*_1_, …, *I*_*N*_ for the source group, and the mixing proportions of the groups *θ* = (*θ*_1_, …, *θ*_*K*_) is defined as

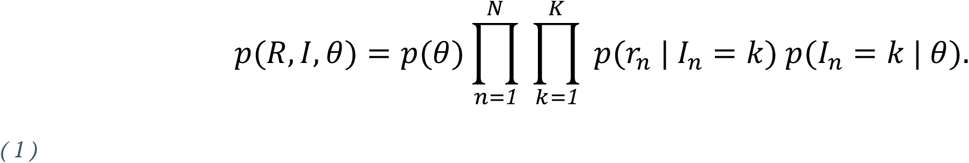

The formulation in Equation (1) corresponds to a mixture model with observations *r*_n_, categorically distributed latent variables *I*_*n*_, event probability parameters *θ*, and the likelihood *p*(*r*_*n*_ | *I*_*n*_ = *k*) of observing the full pseudoalignment count vector *r*_*n*_ given that the group *k* is the true source.

### Likelihood

The likelihood *p*(*r*_*n*_ | *I*_*n*_ = *k*) needs to be defined carefully in order to satisfy the goals of invariance to group identity and size, and monotonicity with increasing pseudoalignment counts within a group. Given the vector *r*_n_, whose components *r*_n,k_ denote the pseudoalignment count in group *k*, we define the likelihood *p*(*r*_*n*_ | *I*_*n*_ = *k*) of observing the whole vector *r*_n_ assuming that *k* is the true group in three parts

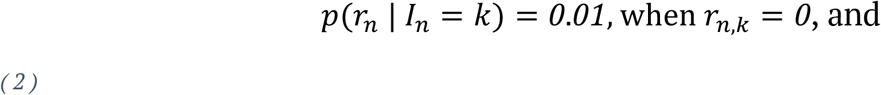

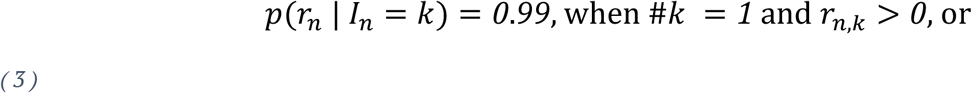

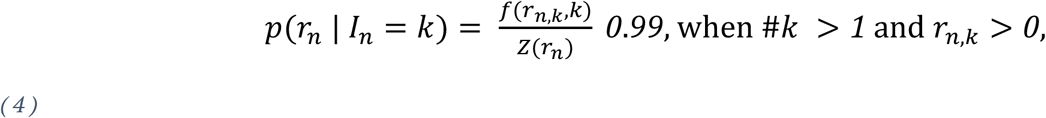

with

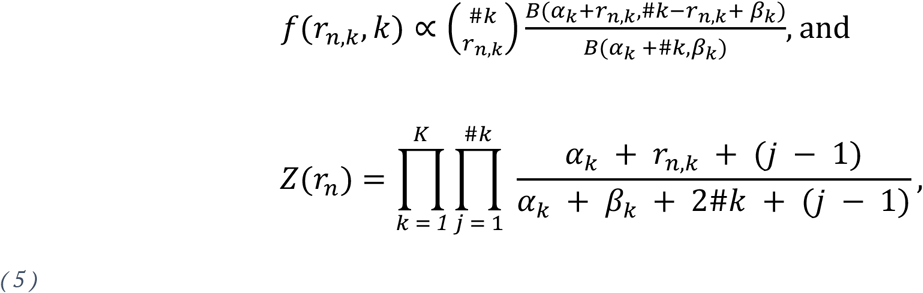

where *B*(*a*, *b*) is the beta function and #*k* is the number of reference sequences in the group *k*. The denominator *Z*(*r*_*n*_) in Equations (4) and (5) arises from deriving the normalizing constant for normalizing *f*(*r*_*n,k*_, *k*) over *r*_n_. The derivation follows from using the identity 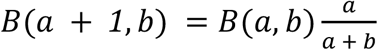 to express each *f*(*r*_*n,k*_, *k*), *k* = {*1*, …, *K*: #*k* > *1*} as a product of the probability mass function of a beta-binomial distribution with parameters (*α*_*k*_ + #*k*, *β*_*k*_), and the term 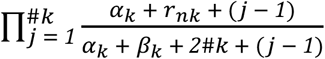 which leads to the form *f*(*r*_*n,k*_, *k*) has in Equation (5). Then, *Z*(*r*_*n*_) is obtained by considering normalizing over the full vector *f*(*r*_*n*_) = (*f*(*r*_*n,1*_, *1*), …, *f*(*r*_*n,k*_, *k*), …, *f*(*r*_*n,K*_, *K*)), where *k* = {*1*, …, *K*: #*k* > *1*}.

The formulation of *f*(*r*_*n,k*_, *k*) in Equations (4) and (5) intuitively arises when the probability mass function of a beta-binomial random variable with hyperparameters (*α*_*k*_, *β*_*k*_) is multiplied by the factor 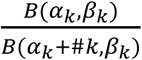. This factor is the inverse of the value of the probability mass function when the observed value is equal to the total number of groups #*k*, meaning in our context a read which is compatible with all reference sequences in a group. Formulating the likelihood in this manner (Equations 4 and 5) causes groups where all sequences in the group are compatible with the read to have equal likelihoods. Compared to a model assuming independence between the groups, this formulation reduces the effect of the likelihoods being flattened in groups with large numbers of assigned reference sequences when compared to small groups, which is necessary to compare groups that differ greatly in size.

Reads with identical pseudoalignment count vectors *r*_n_ have the same likelihood and can be assigned into equivalence classes defined by the count of compatible sequences in each group. This enables a computational optimization where the likelihoods need only be calculated for the observed equivalence classes and then multiplied by the total number of times each equivalence class was observed.

### Model hyperparameters

Instead of using the parametrization (*α*_*k*_, *β*_*k*_) in Equation (5), we use a reparameterization where

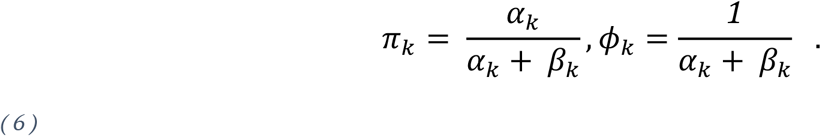

The first parameter *π*_*k*_ corresponds to the mean of the beta distribution that has been compounded with a binomial distribution to obtain the beta-binomial component in Equation 6, and the second parameter *φ*_*k*_ represents a measure of variation in the success probability of each observation [34].

We constrain the mean success rate *π*_*k*_ to *π*_*k*_ ∈ (*0.5, 1*), which produces beta-binomial distributions with an increasing probability mass function [35] in the number of compatible sequences *r*_*n,k*_, which leads to the definition in Equation 5 having the same property. Increasing probability mass functions fulfill our requirement for the likelihood that of two equally sized groups with different number of compatible reference sequences, the one with more compatible sequences is always a better candidate for being the true source. The values of the parameters (*π*_*k*_, *φ*_*k*_) are set to *π*_*k*_ = *0.65*, 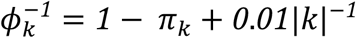 to robustly capture the variance in the alignment count distributions.

### Inference

We perform inference over the mixing proportions *θ* of the different groups using fast collapsed variational inference [36]. The method collapses the mixing proportions *θ* and uses natural gradients to optimise an approximation to the posterior distribution over the indicator variables *I*_*n*_, assuming the distribution factorises over *θ* and *I*_*n*_. The same variational Bayesian method was also used in BitSeqVB [10] to obtain transcript expression levels and has been applied to estimate mixing proportions in bacterial sequencing data in BIB [7] using a different likelihood. The prior distribution on the mixing proportions *θ* is set to Dirichlet (*α*, …, *α*) with *α* = *1*. The same prior was also used by BIB. Since reads originating from the same equivalence class have the same likelihood, variational inference will yield identical posterior inferences for them. This allows us to perform the inference on the smaller number of equivalence classes rather than all reads, leading to faster inference.

### Detection thresholds

Detection thresholds define a means to quantify reliable identification of the reference groups through constructing a minimum relative abundance threshold on the groups. Abundance estimates that fall under the threshold are considered unreliable and set to zero. To obtain the detection thresholds (Figure 1), we generate 100 samples from each reference group within a species by resampling one million sequencing reads, roughly matching the number of reads in our study samples, from the reads used to assemble the reference sequences. Only one reference sequence is used in each new sample. Reads included in the new samples are sampled with replacement with each read having the same probability of being included. Reference sequences used for resampling were chosen at random such that the number of reference sequences from each group corresponds to the square root of the total size of the group. Each group is represented at least once, except for groups which contain only one reference sequence where we apply the maximum detection threshold observed for other groups of the same species. Similarly, species that are represented in the reference by a single group were not resampled from, and the detection threshold was instead fixed at 0.05. After resampling from the reference sequences, the new samples are put through pseudoalignment and mSWEEP abundance estimation without including the reference sequences used in resampling as pseudoalignment targets.

In defining the detection thresholds, the relative abundance estimates obtained from the resampled sequencing reads are represented by 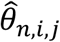, where *n* = 1, …, *N* (in our examples we chose *N* = 100) indicates the new samples resampled from the reference group *i* = 1, …, *K*. The third index *j* = 1, …, *K* denotes the reference group that the abundance estimate was obtained for. We first define source-specific thresholds *q*_*i,j*_ that give a threshold on the reference groups *j* assuming that the true group *i* in the sample is known. The source-specific threshold *q*_*i,j*_ on group *j* ≠ *i* is defined by ordering the relative abundance estimates for the cluster *j*, 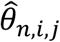, in an ascending order in *n*, and determining the cutoff point *q*_*i,j*_ where *m, m* ∈ {1, …, *N*}, relative abundance estimates fall below the cutoff. Using the source-specific thresholds *q*_*i,j*_, we define the detection threshold *q*_*i*_ on group *i* as *q*_*i*_ = *max*{*max*{*j*: *q*_*i,j*_}, *ε*}, where *ε* is the constant minimum threshold for a specific grouping of the sequences within a species that is observed when comparing the empirical cumulative distribution functions in Supplementary Figure 4. We recommend that *ε* be determined for new species by a synthetic mixing procedure similar to what was used to compare the accuracy of mixture estimates to their single-colony counterparts in Figure 4.

Based on the selected value of *m*, we define a statistical confidence score for the abundance estimates exceeding the detection threshold *q*_*i*_ as

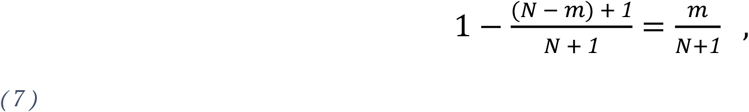

which corresponds roughly to the fraction of resampled samples that exceed the threshold corresponding to the value of *m*. Using a value of *m* closer to the number of samples in constructing the detection thresholds result in stricter thresholds and thus more confidence in the abundance estimates exceeding the threshold. Results reported in our experiments include thresholds constructed with *m* = 100 and *m* = 90, corresponding to confidence scores (Equation 7) of approximately 0.99 and 0.90, respectively.

### *E. coli* plate sweeps from Vietnamese children

In Ho Chi Minh City, 750 children were recruited into a diarrhoeal cohort study and followed for 2 years. Stool samples were collected at routine sampling points and when the children had an episode of diarrhoea. All stool samples were cultured to identify pathogens and onto MacConkey plates to isolate *E. coli* and other *Enterobacteriaceae*. The MacConkey plates were scraped and stored in 20% glycerol at −80°C. The frozen plate sweeps from 48 diarrhoea episodes, paired with 48 asymptomatic samples (96 in total), were revived on MacConkey media; plates were scraped and total genomic DNA was extracted using the Wizard genomic DNA purification kit (Promega, USA). The extracted DNA was sequenced using the Illumina HiSeq platform using the method described elsewhere [37]. Antimicrobial resistance genes were detected using the ARIBA software [19]. The raw sequence data data can be found in the ENA under the accession numbers detailed in table Supplementary Table 5.

## Declarations

### Ethics approval and consent to participate

Ethical approval was required for the cohort study contributing the *E. coli* organisms. Approvals were provided by the Oxford University Tropical Research Ethics Committee (OxTREC approval 1058–13) as well as from local partners including the Institutional Review Board (IRB) at the Hospital for Tropical Diseases (HTD) and Hung Vuong Hospital (HVH). Written informed consent was obtained from the parent or guardian of all children.

### Availability of data and material

mSWEEP software is available in GitHub: https://github.com/PROBIC/mSWEEP. Reference sequences and the grouping used in producing the results have been compiled and submitted to figshare under the doi 10.6084/m9.figshare.8222636. Accession numbers for the reference data can be found in Supplementary Table 4. Accession numbers for the 96 Vietnamese *E. coli* plate sweeps are available in Supplementary Table 5. Accession numbers for the *K. pneumoniae* mixture samples and 39 *Campylobacter* mixture samples are available in Supplementary Table 6. The remaining 77 *Campylobacter* mixture samples have been submitted to figshare under DOIs 10.6084/m9.figshare.6445136 and 10.6084/m9.figshare.6445190.

### Author Contributions

TM, JC, and AH developed the model and designed the method comparison experiments. TM implemented the model and ran the mSWEEP analyses. TK built the *K. pneumoniae* and *E. coli* references. CJB, SB, and JP collected the *E. coli* data from Vietnam, ran the antimicrobial resistance analyses, and interpreted all results from the *E. coli* data. SD, CJB, BP, GM, DMA, EJF, and SKS collected the *K. pneumoniae* and *C. jejuni* mixture data and interpreted the results from it. TM, TK, SD, CJB, SKS, JC, and AH wrote the paper. JC and AH conceived the study. All authors discussed the results and commented on the manuscript.

