## Supplementary Figures 1-7 for "High-resolution sweep metagenomics using fast probabilistic inference"

Lineage abundances in 129 *K. pneumoniae* single−colony samples

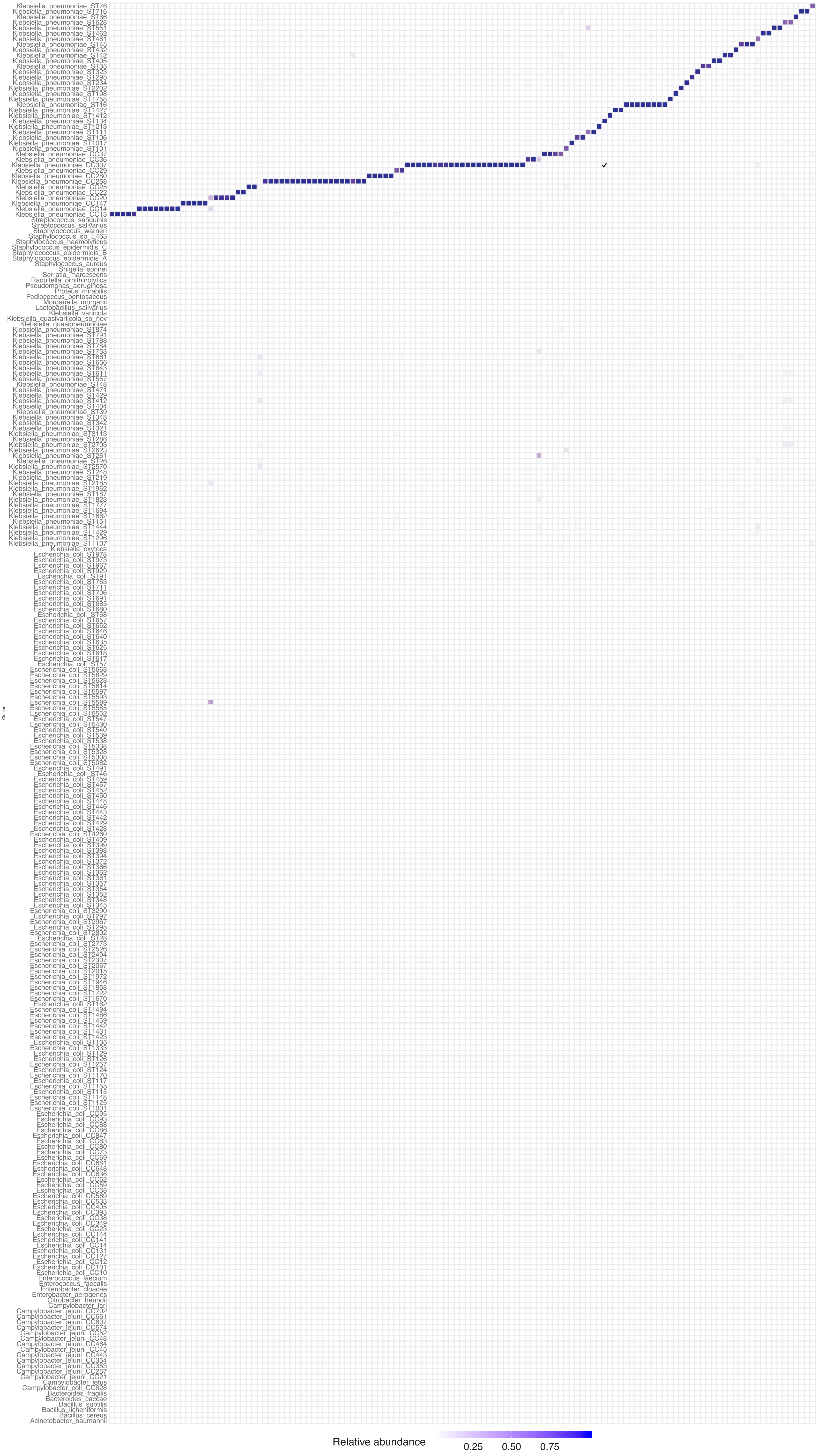


Supplementary figure 1a Lineage abundances in 129 K. pneumoniae single−colony samples. Visualization of the relative abundance estimates. The diagonal contains the true cluster of origin in each sample while deviations from the diagonal indicate errors in true negative groups. The three incorrectly identified samples are visible as columns where the abundance estimates are spread across multiple cells.

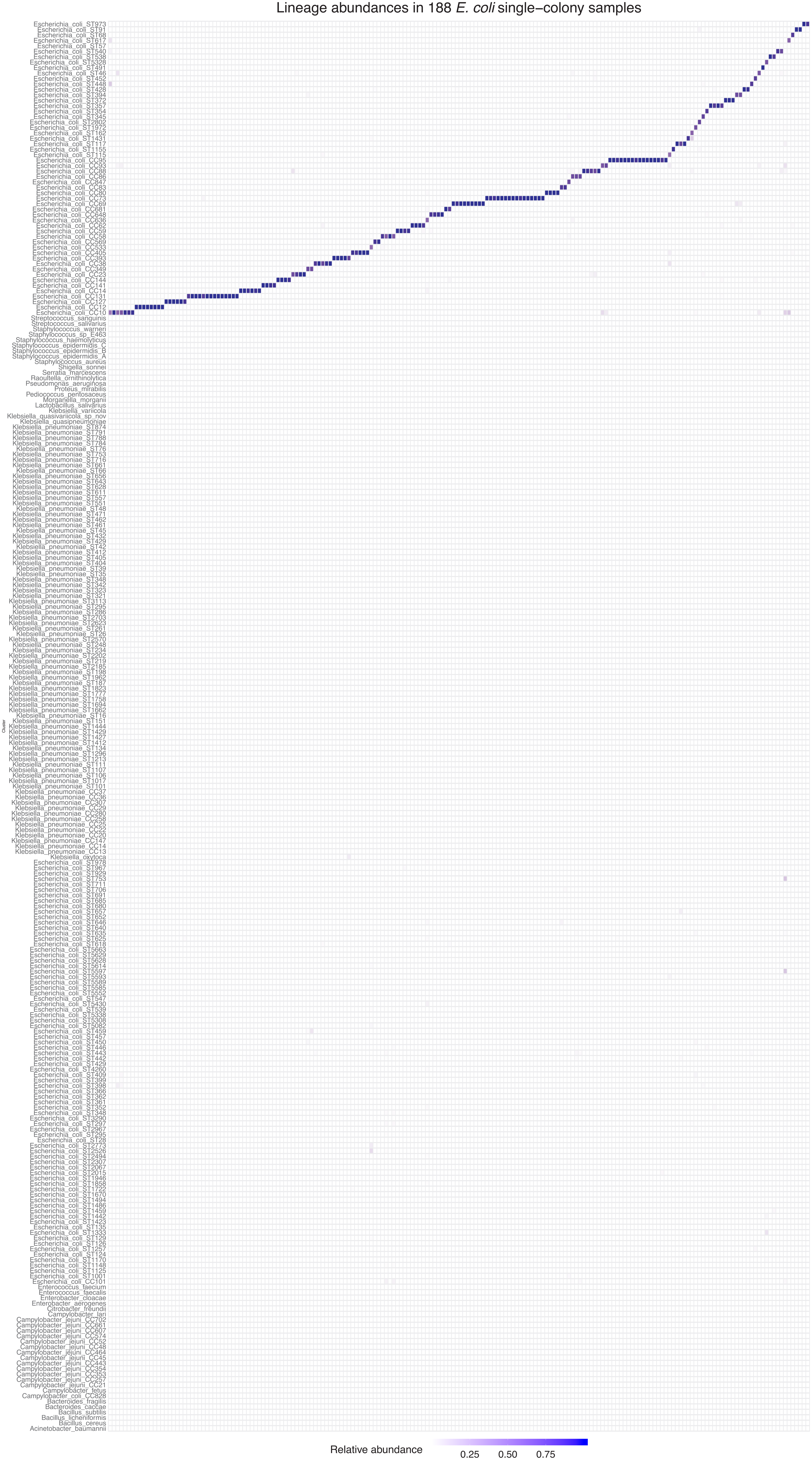


Supplementary figure 1b Lineage abundances in 188 E. coli single−colony samples. Visualization of the relative abundance estimates. The diagonal contains the true cluster of origin in each sample while deviations from the diagonal indicate errors in true negative groups. The three incorrectly identified samples are visible as columns where the abundance estimates are spread across multiple cells.

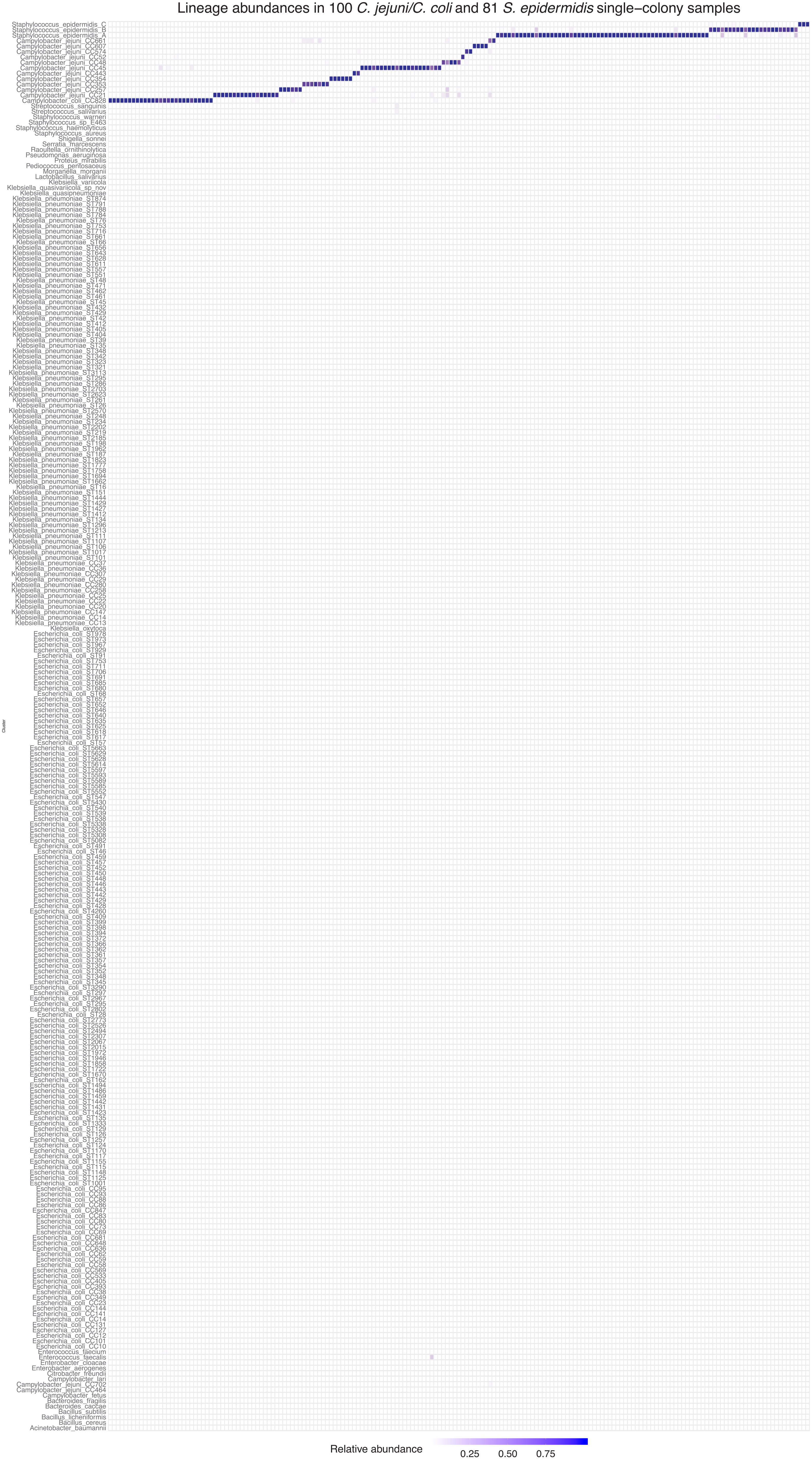


Supplementary figure 1c Lineage abundances in 100 C. jejuni/C. coli and 81 S. epidermidis single−colony samples. Visualization of the relative abundance estimates. The diagonal contains the true cluster of origin in each sample while deviations from the diagonal indicate errors in true negative groups. The three incorrectly identified samples are visible as columns where the abundance estimates are spread across multiple cells.

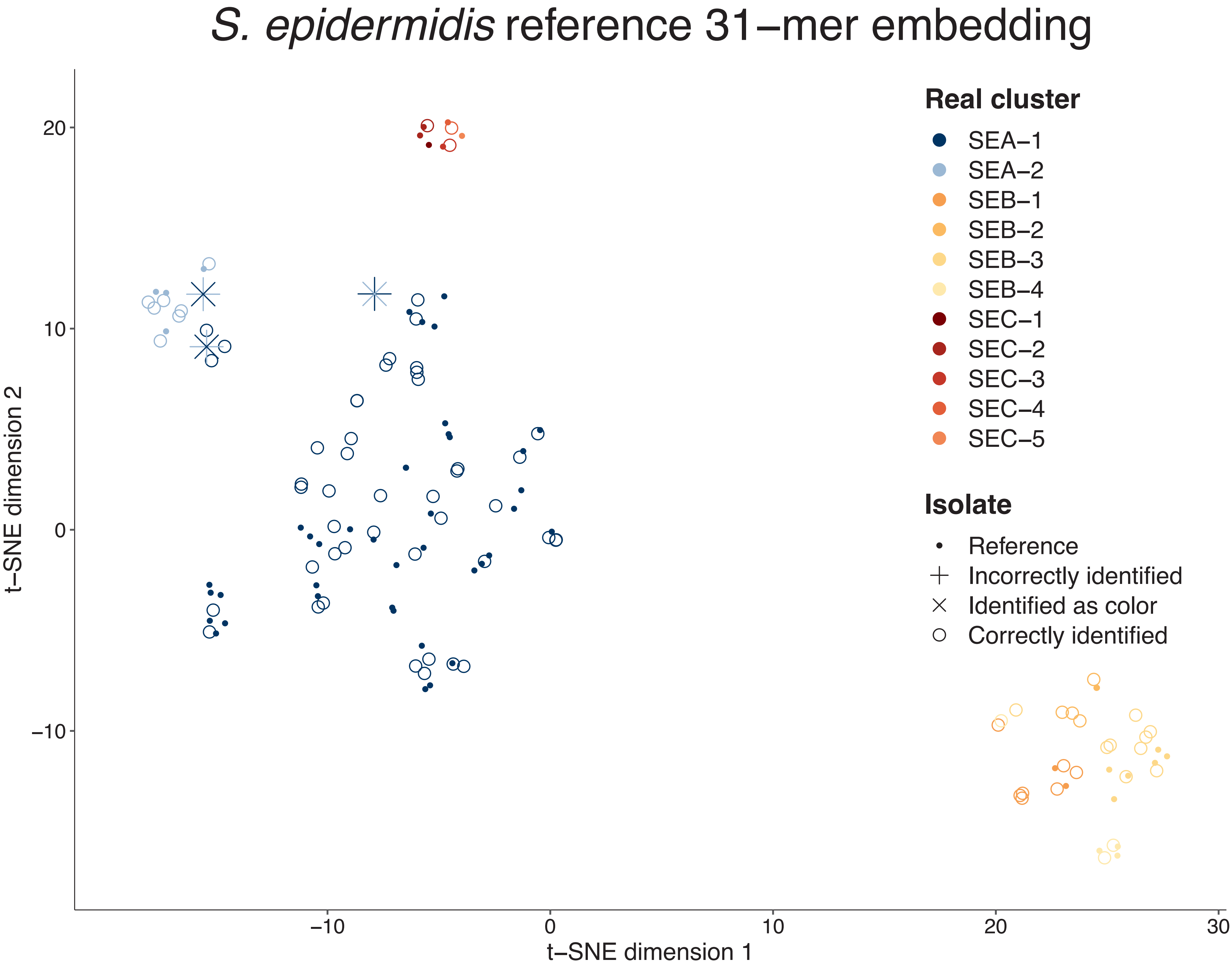


Supplementary figure 2a S. epidermidis reference 31−mer t−SNE embedding of the reference sequences. The first level of BAPS−clustering (SEA, SEB, and SEC) is well separated in placement of the sequences. The further split into subgroups of SEA, SEB, and SEC is less clear with sequences from the subgroups being mixed with each other. The uncertainty in the grouping results in inaccuracies in relative abundance estimation.

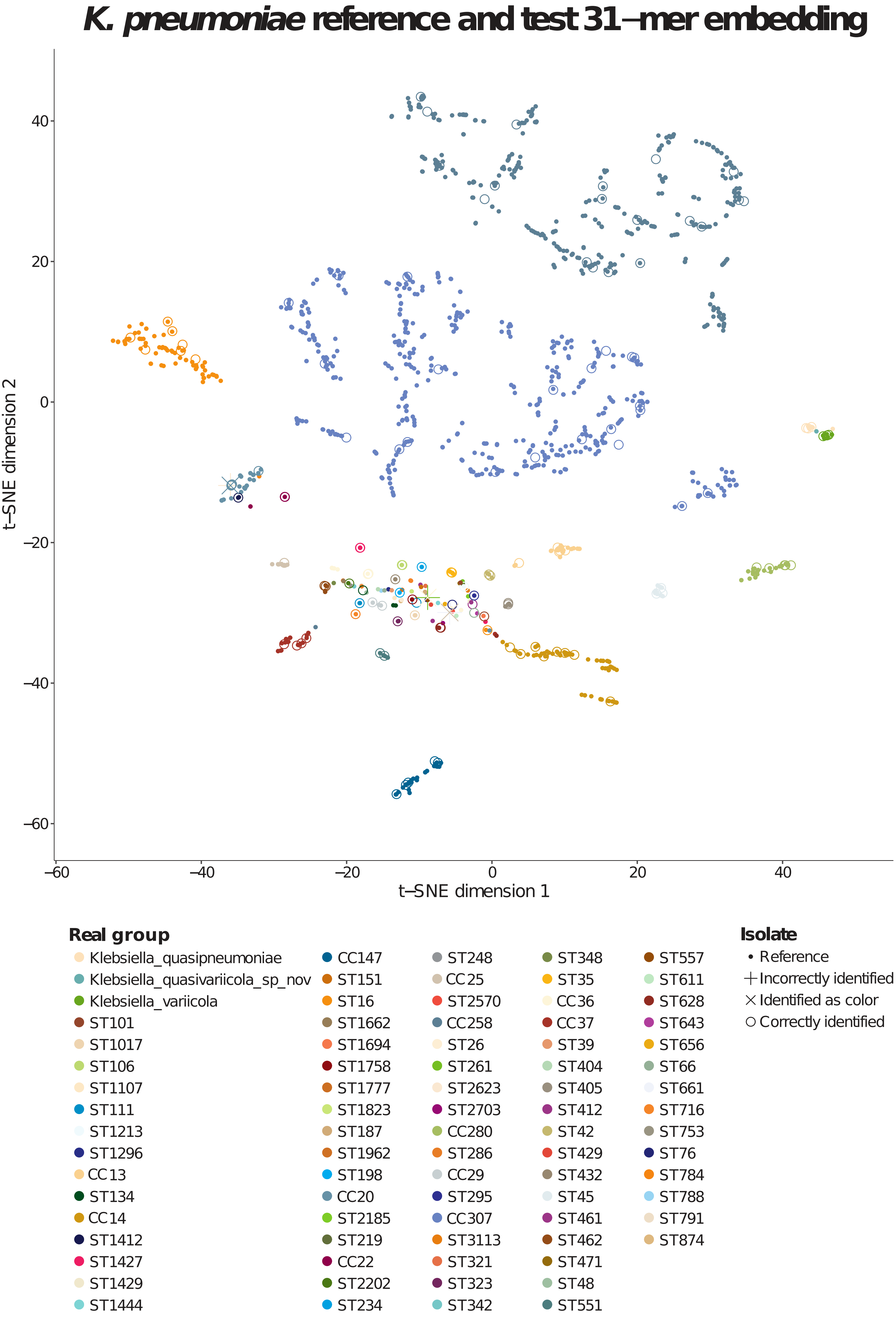


Supplementary figure 2b K. pneumoniae reference 31-mer embedding. The t-SNE embedding for the K. pneumoniae reference contains multiple well-defined larger groups along with a wider variety of single-sequence groups.

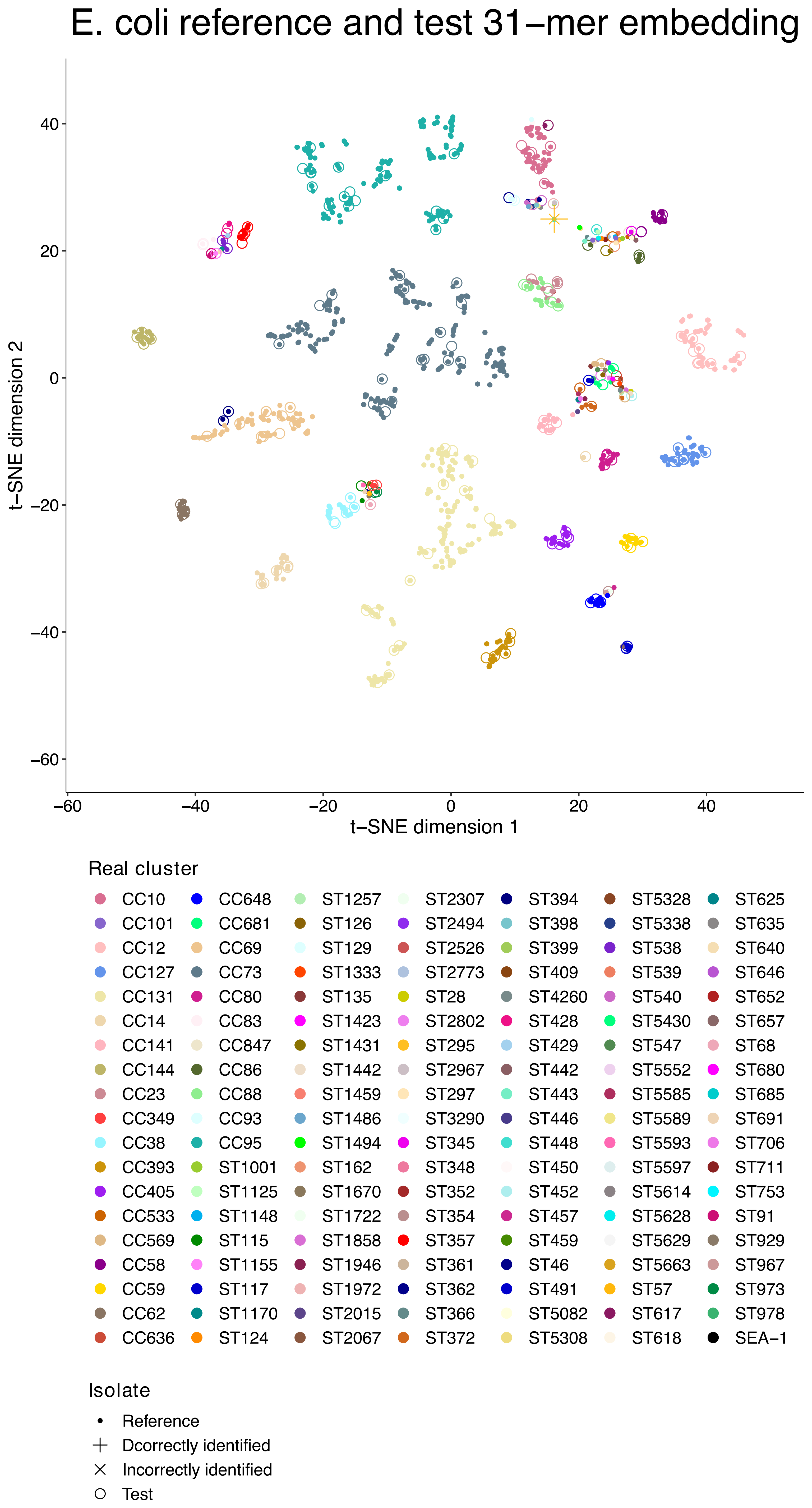


Supplementary figure 2c E. coli reference 31−mer t−SNE embedding of the reference sequences. The E. coli references conform well to the sequence type and clonal complex groupings. Groups represented by single sequences are placed in their own groups rather than within the established lineages, while the lineages containing multiple sequences form distinct clusters.

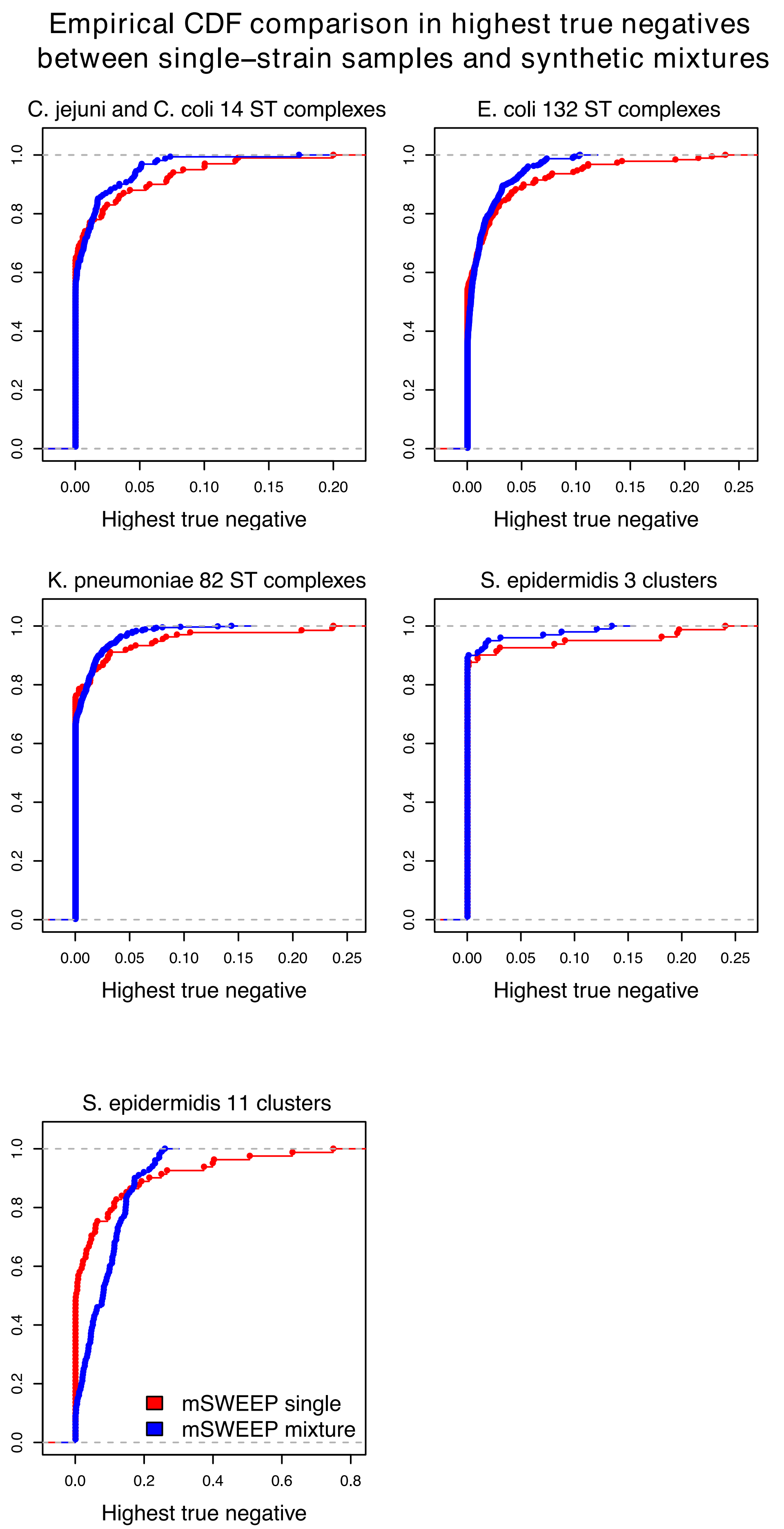


Supplementary figure 3 Empirical CDF comparison in highest false positives between single-strain samples and synthetic mixtures. Except the S. epidermidis 11 cluster case, the error distributions obtained from synthetic mixtures when measured by the false positive estimates are nearly identical or contain smaller errors than what is obtained from single-strain isolates. The comparison demonstrates that if the single-strains are accurately identified, mixtures containing multiple strain groups are not more difficult to identify.

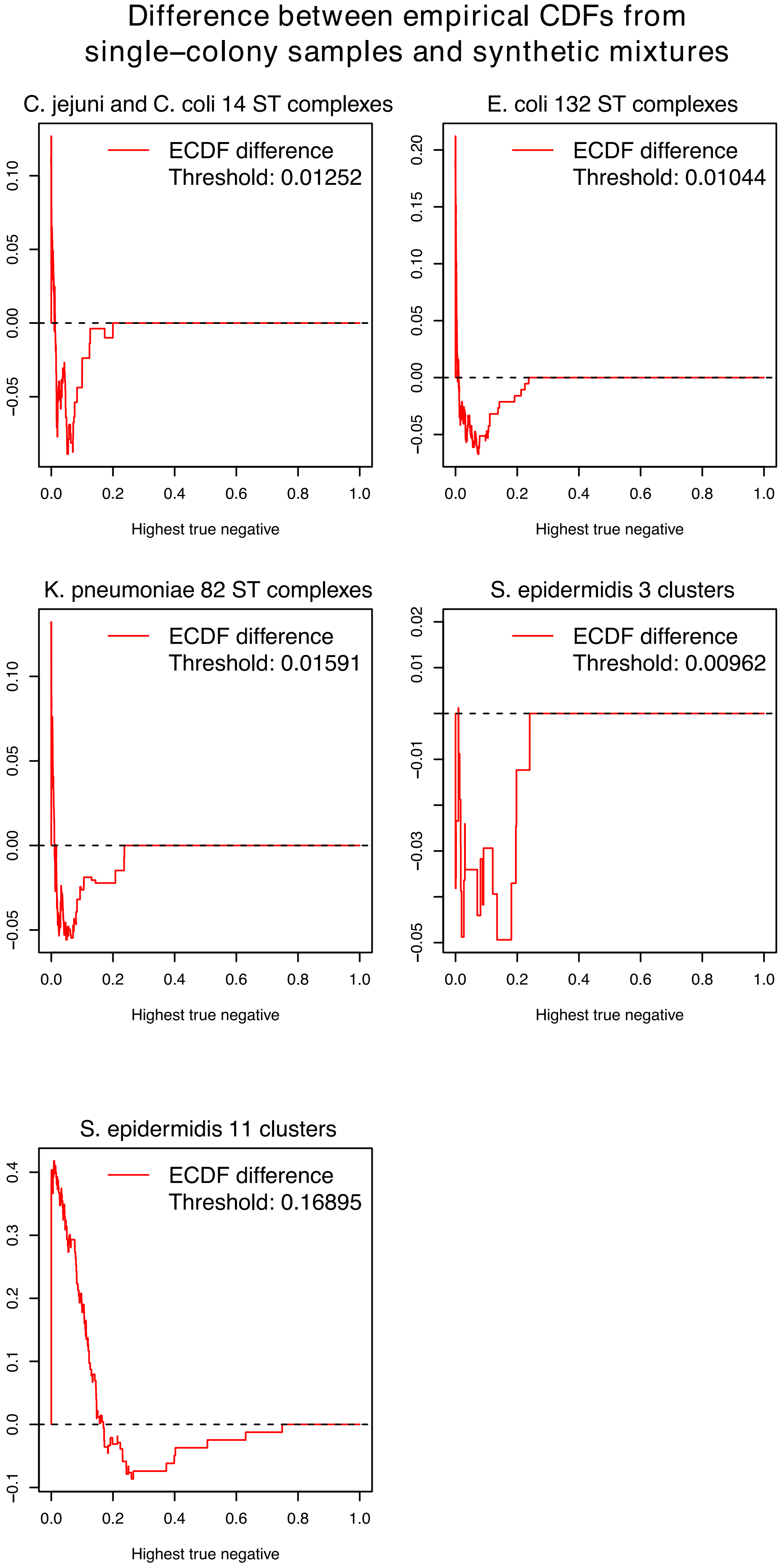


Supplementary figure 4 Difference between empirical CDFs from single-strain samples and synthetic mixtures. Except the S. epidermidis 11-cluster case, the difference between the ECDFs from Supplementary Figure 4 shows that the mixture samples produce more accurate estimates after exceeding the threshold of 0.016, and that the area above the single-strain ECDF but under the mixture ECDF exceeds the area above the mixture ECDF but below the single-strain curve.

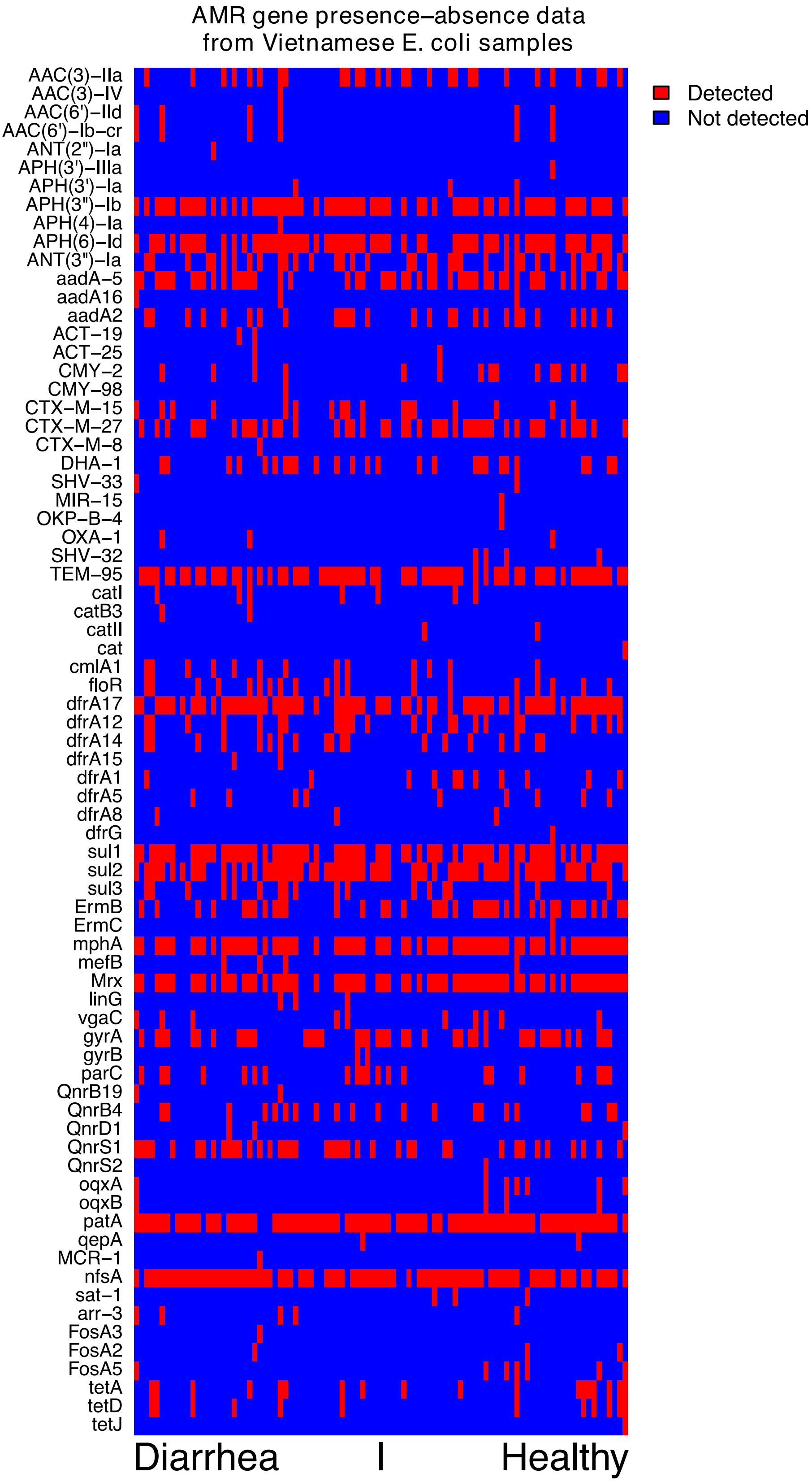


Supplementary figure 5 Antimicrobial resistance (AMR) gene presence-absence in Vietnamese E. coli samples. The genes (rows) were detected in the samples (columns) using the ARIBA software. Red denotes the detection of a AMR gene in a sample, and blue absence. Diarrheal samples are shown on the left half of the plot, and asymptomatic samples on the right half.

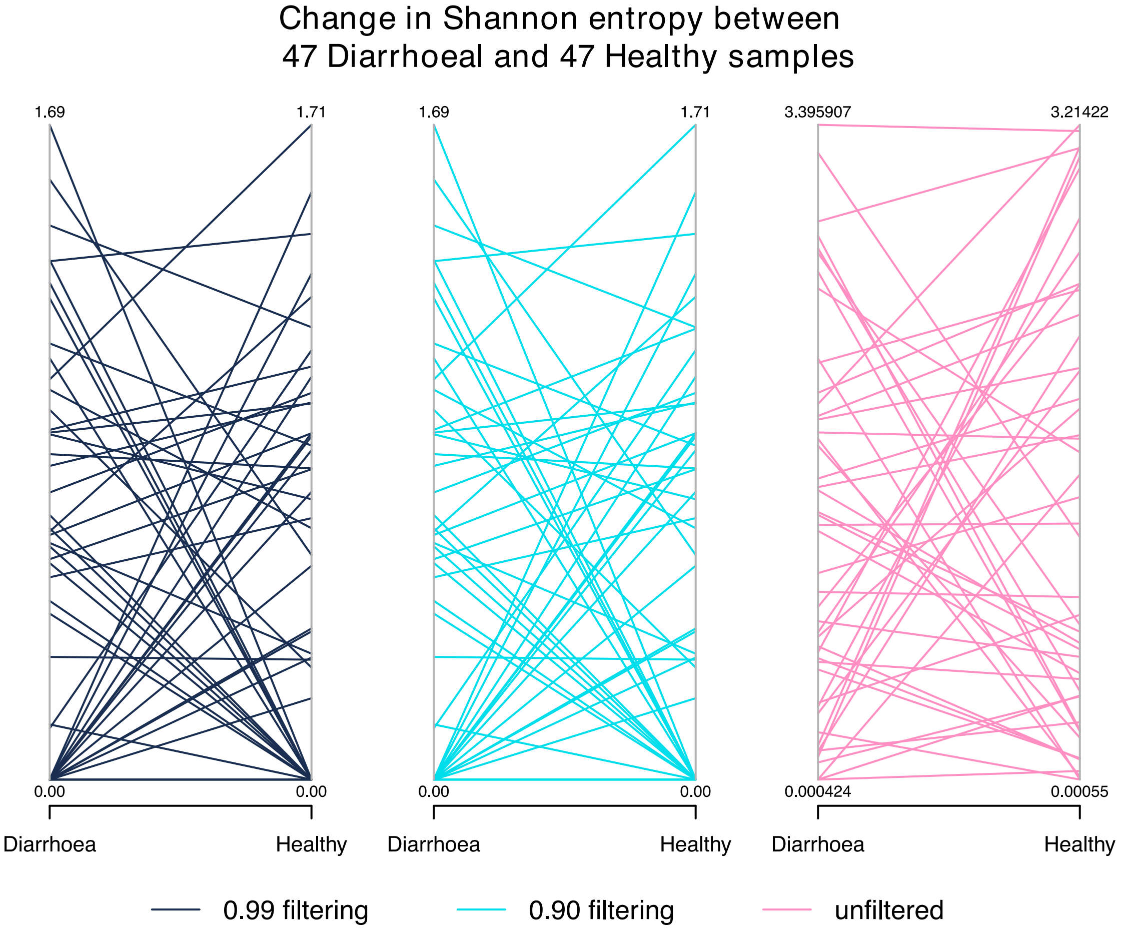


Supplementary figure 6 Change in Shannon entropy between diarrhoeal and asymptomatic E. coli samples from Vietnamese children. There are no major changes in the overall distribution of the Shannon entropies between the two sampling points.

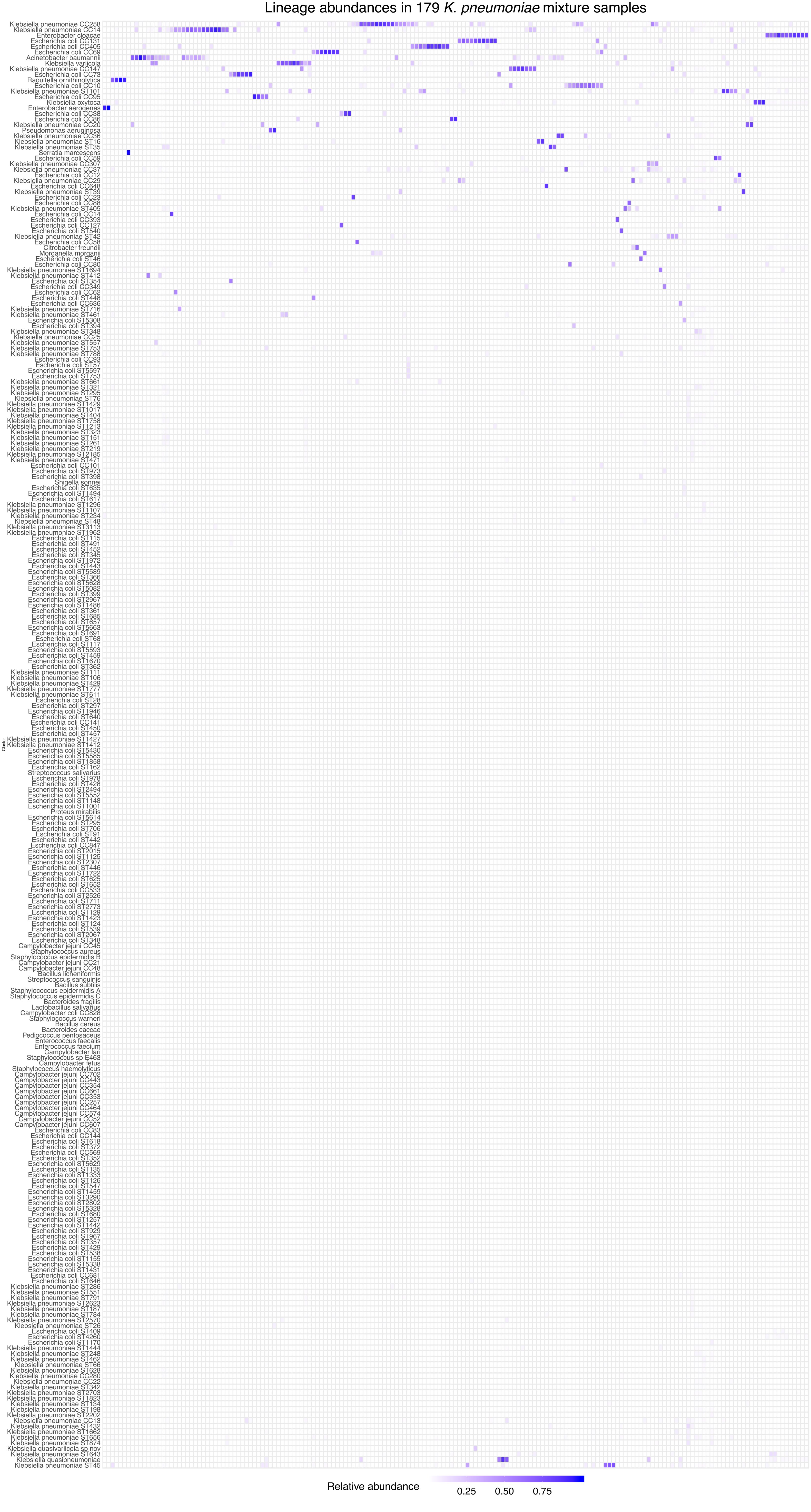


Supplementary figure 7 Lineage abundances in 179 K. pneumoniae mixture samples. Relative abundance estimates originating from mixture samples containing E. coli and K. pneumoniae visualized as a heatmap. Most of the samples contain sequencing data from multiple groups.
